# Complete sequences of *Schizosaccharomyces pombe* subtelomeres reveal multiple patterns of genome variation

**DOI:** 10.1101/2020.03.09.983726

**Authors:** Takuto Kaji, Yusuke Oizumi, Sanki Tashiro, Yumiko Takeshita, Junko Kanoh

## Abstract

Genome sequences have been determined for many model organisms; however, repetitive regions such as centromeres, telomeres, and subtelomeres have not yet been sequenced completely. Here, we report the complete sequences of subtelomeric homologous (*SH*) regions of the fission yeast *Schizosaccharomyces pombe*. We overcame technical difficulties to obtain subtelomeric repetitive sequences by constructing strains that possess single *SH* regions. Whole sequences of *SH* regions revealed that each *SH* region consists of two distinct parts: the telomere-proximal part with mosaics of multiple common segments showing high variation among subtelomeres and strains, and the telomere-distal part showing high sequence similarity among subtelomeres with some insertions and deletions. The newly sequenced *SH* regions showed differences in nucleotide sequences and common segment composition compared to those in the *S. pombe* genome database (PomBase), which is in striking contrast to the regions outside of *SH*, where mutations are rarely detected. Furthermore, we identified new subtelomeric RecQ-type helicase genes, *tlh3* and *tlh4*, which add to the already known *tlh1* and *tlh2*, and found that the *tlh1–4* genes show high sequence variation. Our results indicate that *SH* sequences are highly polymorphic and hot spots for genome variation. These features of subtelomeres may have contributed to genome diversity and, conversely, various diseases.

## Introduction

Genomic DNA sequences provide fundamental information for biological study. The genomes of model organisms, such as *Saccharomyces cerevisiae* (*S. cerevisiae*) (1), *Schizosaccharomyces pombe* (*S. pombe*) (2), *Ceanorhabditis elegans* (3), *Drosophila melanogaster* (4), *Arabidopsus thaliana* (5), and *Homo sapiens* (6–8), have been sequenced in the last two decades, and most parts of these sequences have been reported. However, sequencing of long repetitive regions has not been completed because of technical difficulties in sequencing and chromosome allocation as well as relatively frequent mutations and structural changes caused by chromosome rearrangements such as recombination, translocation, chromosome breakage, and fusion in these regions (9–13).

Incomplete genomic DNA information can lead to inaccurate data in some experiments. For instance, we are unable to determine the precise chromatin localization of proteins in repetitive regions without actual DNA sequences. Evaluation of protein localization by chromatin immunoprecipitation (ChIP) assays involves PCR with sets of representative primers that target repetitive sequences or Southern blot analysis using representative probes. Chromatin localization values obtained using representative primers or probes are only averages at regions with representative sequences, but they do not reflect actual patterns of chromatin association. This problem is not fully solved by next-generation sequencers without complete genome sequences. In addition, uncharacterized genes may be present in un-sequenced regions. Therefore, complete sequences of genomic DNA are crucial for accurate analyses and a deeper understanding of model organisms.

Telomeres, which exist at chromosome ends and possess species-specific tandem repeat sequences, play crucial roles in several cellular activities required for cell survival, including protection of chromosome ends, length regulation of telomere-specific repeat DNA, and regulation of chromosome movements during mitosis and meiosis (14–17). Subtelomeres, which are adjacent to telomeres, have sequences distinct from telomere repeats and generally contain multiple species-specific segments that share high similarity with other subtelomeres. In the budding yeast *S. cerevisiae*, the subtelomeres contain X and Y’ elements, the latter of which includes the open reading frame (ORF) of a helicase gene (18). In humans, the subtelomeres are mosaics of approximately 50 types of multiple segments containing various ORFs such as those for the *DUX4* gene, which is related to facioscapulohumeral muscular dystrophy, and for the olfactory receptor family genes (9,19,20). Although substantial knowledge of telomeres has accumulated, research on subtelomeres has progressed slowly compared with research on other chromosomal regions with complete DNA sequences because of technical difficulties caused by long, repetitive and partially unknown sequences.

The fission yeast *S. pombe* is one of the most commonly used yeast model organisms for biological study. It usually proliferates as haploid in nutrient-rich media and possesses only three chromosomes (chromosome 1 [Ch1], 5.6 Mb; Ch2, 4.6 Mb; Ch3, 3.5 Mb), which enables the whole package of genetic analyses such as screening for both dominant and recessive mutations, as well as generation of cells with circular chromosomes by deleting telomere DNA. *S. pombe* subtelomeres spanning ∼100 kb are sub-divided into two regions of ∼50 kb each, the telomere-adjacent and telomere-distal regions. The telomere-adjacent regions of subtelomeres contain subtelomeric homologous (*SH*) sequences, which share high similarity (> 98%) with at least one other subtelomere in *S. pombe* (21) and form heterochromatin structures (22,23). In contrast, the telomere-distal regions share almost no sequence similarity with other subtelomeres but exhibit common highly condensed chromatin structures called knobs (24) (Figure 1A). Due to high sequence similarity, it is very difficult to distinguish individual *SH* regions of subtelomeres at different chromosome ends. Therefore, parts of *SH* regions remain un-sequenced 18 years after most parts of the *S. pombe* genome sequence were reported (PomBase: http://www.PomBase.org/status/sequencing-status) (2) (Figure 1B). Previously, parts of the four *SH* regions have been cloned and sequenced (pNSU series, see below) (25); however, they have not yet been allocated to specific subtelomeres (PomBase).

**Figure 1.**
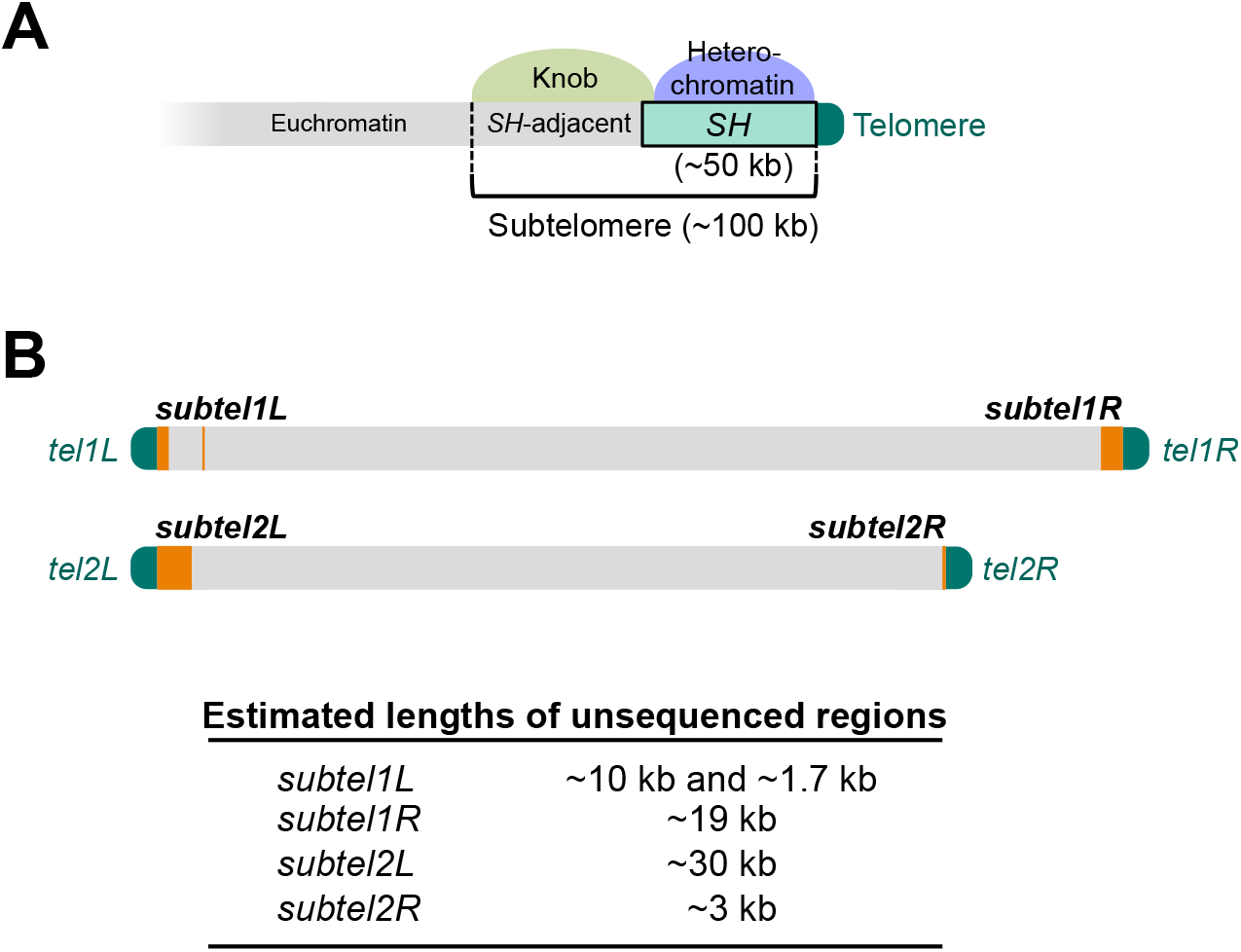
Structures and previously un-sequenced regions of subtelomeres in *S. pombe*. (A) Schematic illustration of the structures of subtelomeres (∼100 kb) of chromosomes 1 and 2 in *S. pombe*. The telomere-adjacent region (∼50 kb) contains an *SH* sequence, which shows high identity (>98%) with sequences of other subtelomeres. Subtelomeric heterochromatin is formed around the *SH* region (21). In contrast, the *SH*-adjacent region (∼50 kb) shows low sequence identities with other subtelomeres but forms a highly condensed knob structure that is shared between them (24). (B) Schematic illustration of un-sequenced regions of subtelomeres in chromosomes 1 and 2 of strain *972* according to PomBase (indicated by orange boxes). Lengths of un-sequenced regions are estimated based on the assumption that these *SH* sequences show high similarity with that of *subtel2R* of PomBase. *Tel1L*, *tel1R*, *tel2L*, and *tel2R* indicate telomeres at the left and right arms of Ch1 and those of Ch2, respectively. *Subtel1L*, *subtel1R*, *subtel2L*, and *subtel2R* indicate subtelomeres at the left and right arms of Ch1 and those of Ch2, respectively.

## Materials and Methods

### Strains and general techniques for *S. pombe*

*S. pombe* strains used in this study are listed in Supplementary Table S1. Growth media and basic genetic and biochemical techniques used in this study were described previously (26–28).

### Construction of *972SD4* strains

To construct *972SD4* strains containing single *SH* regions of the standard wild-type strain *972*, the *SD5* strain (ST3524) (21), in which all five *SH* regions were replaced with selective marker genes (*his7*^+^ or *ura4*^+^), was crossed with strain *972*, and the progeny were crossed with the *SD5* strain (ST3479 or ST3524) again. The presence or absence of each *SH* region in the resulting progeny were examined by pulse-field gel electrophoresis (PFGE) followed by Southern blotting.

### PFGE

PFGE of NotI-digested chromosomal DNA was performed using a CHEF-DR III Pulsed Field Electrophoresis System (BioRad) under the following conditions: 1% SeaKem Gold Agarose (Lonza) in 0.5×TBE; temperature, 10°C; initial switch time, 40 s; final switch time, 80 s; run time, 18 h; voltage gradient, 6.8 V/cm; and angle, 120°.

### Southern blotting

NotI-digested chromosomal DNA was separated by PFGE and subjected to Southern blotting. For the telomere and telomere-associated sequence (TAS) probes, telomeric DNA and TAS fragments (TAS1, TAS2, and TAS3) were excised from pNSU70 (25). For the probe that specifically recognizes the *SH* regions of subtelomeres in the left arm of Ch1 (*subtel1L*) and the left and right arms of Ch2 (*subtel2L* and *subtel2R*) but not the subtelomere of the right arm of Ch1 (*subtel1R*), the *SPBCPT2R1.03* ORF was amplified by PCR. These DNA fragments were labeled with digoxigenin (DIG) using a DIG High Prime DNA Labeling and Detection Starter Kit II (Roche), and signals were detected according to the manufacturer’s instructions.

### Cloning and sequencing of *SH* regions

DNA fragments containing telomere-proximal *SH* regions (∼5 kb) were amplified by PCR from genomic DNA of each *972SD4* strain using Phusion High-Fidelity DNA polymerase (Thermo Fisher) and the following primer set.

jk1861: 5’-ACTAGTGGATCCCCCTGTAACCACGTAACCTTGTAACC-3’

jk1862: 5’-GAATTCCTGCAGCCCGGTTTGAGCATCTGTCAGAGGTAA-3’

Each DNA fragment was inserted at the Sma1 site of pBluescript SK(–) (Stratagene) using In-Fusion HD Cloning Kit (Clontech). The resulting plasmids were digested with Kpn1 and Xho1 at the multiple cloning site of the vector and serially deleted from the XhoI cutting site (5’-protruding end) by treating the plasmids with exonuclease III and mung bean nuclease (Takara) for fixed times. After both ends of the plasmids were blunted using Klenow fragments (Takara), they were ligated by DNA ligase (Takara).

Each re-circularized plasmid was cloned using *Eschelicia coli* (*E. coli*), XL1-Blue (*recA1 endA1 gyrA96 thi-1 hsdR17 supE44 relA1 lac* [*F’proAB lacI^q^ZΔM15 Tn10* (Tet^r^)]), and then the DNA was sequenced using the following primers that anneal to the vector.

st13 (M13 primer M3): 5’-GTAAAACGACGGCCAGT-3’

st14 (M13 primer RV): 5’-CAGGAAACAGCTATGAC-3’

Portions of telomere-proximal *SH* sequences were assembled using their overlapping sequences (Supplementary Figure S1). Two independent strains of each *972SD4* were analyzed.

To sequence most of the telomere-distal non-repetitive *SH* regions, DNA fragments (1.3–2.9 kb) were amplified by PCR and the sequences were determined, whereas sequences of regions with some repeats were determined using the deletion method described above. Two independent strains of each *972SD4* were analyzed.

DNA sequences were determined using BigDye Terminator v3.1 Cycle Sequencing Kit (Applied Biosystems), Prism 3130*xl* Genetic Analyzer (Applied Biosystems), and DNA Sequencing Analysis Software v5.4 (Applied Biosystems).

### Classification of telomere-proximal *SH* sequences into common segments

First, the sequences of telomere-proximal *SH* regions were classified into common segments that meet the criteria of ≥50 bp and >95% identity using NCBI BLAST program (https://blast.ncbi.nlm.nih.gov/Blast.cgi). Second, gaps between the segments were classified into additional segments that meet the criteria ≥14 bp and >95% identity.

To minimize the number of common segments, we set exceptional rules for subtypes of C, E, K, and S, which contain different copies of common sequence motifs that show 100% identities except for motifs in subtype C, c1–8 (Figure 3B). Segments were classified into variants that meet the criteria of 100% identity (Figure 3A and B, and Supplementary Figure S2).

### RNA analyses

Total RNA was purified from exponentially growing cells as described previously (29). For the reverse transcription (RT)-PCR, complementary DNA was synthesized using a High-Capacity cDNA Reverse Transcription Kit (Applied Biosystems) with random primers and analyzed by conventional PCR (Figure 6C) or quantitative PCR (Figure 6D) using a StepOne Real-Time PCR System. Primer sequences are listed in Supplementary Table S2.

## Results

### Construction of strains containing single *SH* regions of *972*

To overcome the difficulty in allocating each *SH* sequence to a specific subtelomere, we constructed strains containing single *SH* regions of the standard wild-type strain *972*. Strain *972* (*h*^−^) used in this study (a derivative of the original *972* (30)), which has not been crossed with other strains, possesses four *SH* sequences (*SH1L*, *SH1R*, *SH2L*, and *SH2R*, as shown in Figure 2A) adjacent to the telomeres of Ch1 and Ch2, but no *SH* sequences in Ch3 (note that some strains also possess a partial (∼15 kb-long) *SH* sequence adjacent to the telomeres of the left and/or right arms of Ch3) (21,29,31). We previously produced the *SD5* (<u>s</u>ubtelomere <u>d</u>eletion 5) strain in which all five *SH* regions that are located at both ends of Ch1 and Ch2 and the left end of Ch3 in a non-standard strain JP1225 (a derivative of CHP429 [*h*^−^ *ade6-M216 leu1-32 ura4-D18 his7-366*] from C. Hoffman) were replaced with a marker gene (*his7*^+^ or *ura4*^+^) (21). Strain *972* was crossed with *SD5*, and the progeny were analyzed for the presence or absence of each *SH* region by PFGE followed by Southern blotting (Figure 2A-D). We screened for strains that exhibit a single band of TAS (i.e., *SH*). We obtained strains that contain a single *SH* region of *972* and named them *972SD4[1L*+, *1R*+, *2L*+, and *2R*+] since they carry deletions of four *SH* regions in the original *SD5* and one intact *SH* region from *972*. Each intact *SH* region in *972SD4* was named as *972SD4-SH1L*, *972SD4-SH1R*, *972SD4-SH2L* or *972SD4-SH2R*.

**Figure 2.**
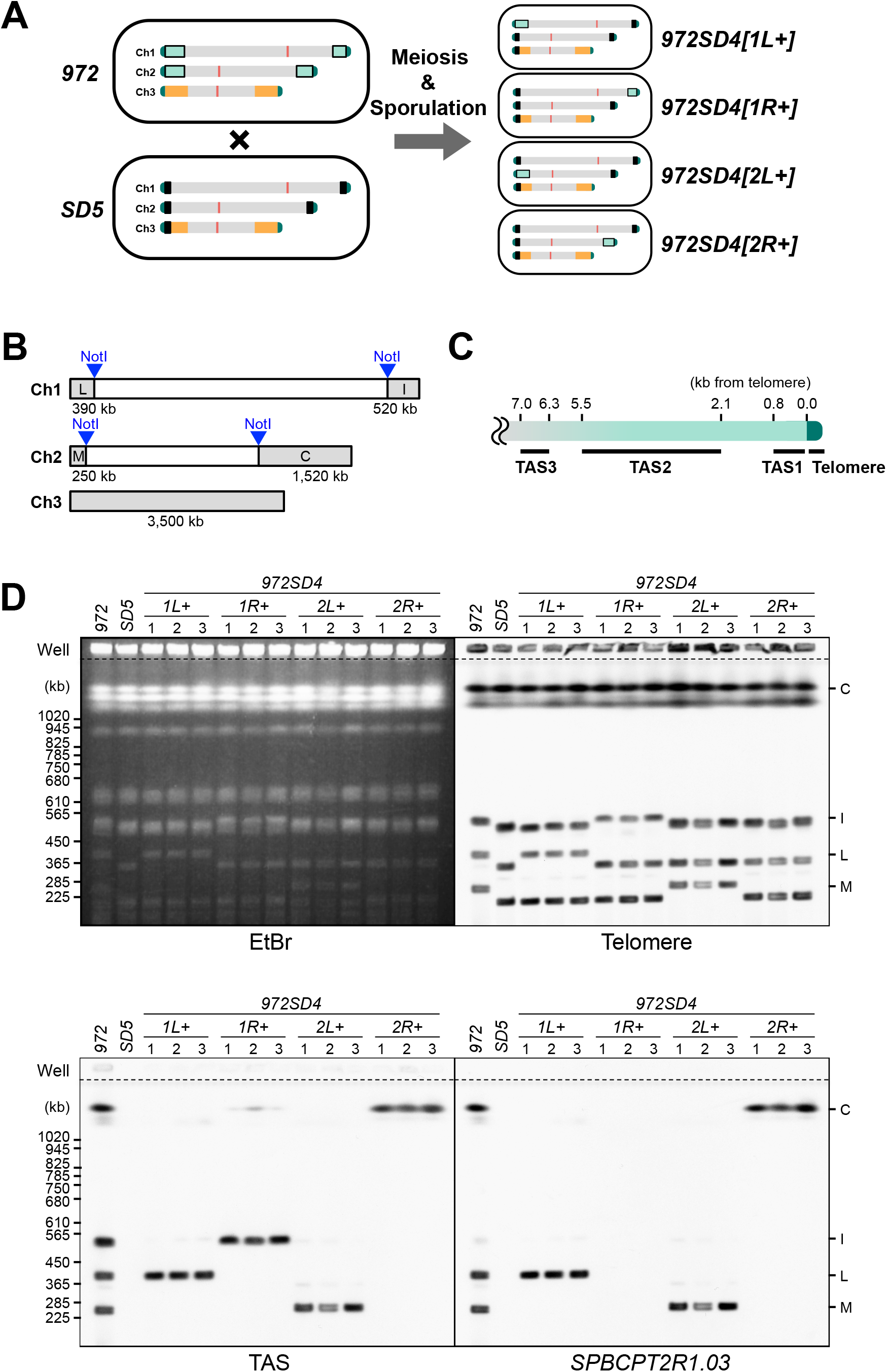
Construction of *972SD4* strains containing a single *SH* region of *972*. (A) Construct strategy of *972SD4* strains. Pale green boxes, *SH* regions adjacent to telomeres; black boxes, replacement of *SH* regions with marker genes (*his7*^+^ or *ura4*^+^) (21); yellow boxes, rDNA repeats; and red boxes, centromeres. Strains *972* and *SD5* were crossed, and meiosis and sporulation (spore formation) were induced. Progenies with single *SH* regions were obtained. (B) Schematic illustration of telomere-containing NotI restriction fragments. Fragments L, I, M, and C contain *SH1L*, *SH1R*, *SH2L*, and *SH2R*, respectively. (C) Schematic illustration of the positions of DNA fragments detected by probes for telomeres and TAS (telomere-associated sequence, TAS1–3). Distances from telomeres in pNSU70 are shown. (D) Analyses of the chromosome end structures in *972SD4* strains. NotI-digested chromosomal DNAs of three independent strains of each *972SD4* were analyzed by PFGE followed by Southern blotting using telomere, TAS, or *SPBCPT2R1.03* ORF probes. EtBr, ethidium bromide staining of the gel after PFGE. Note that *SH1R* lacks part of the *SH* region homologous to *SPBCPT2R1.03* according to PomBase. Consistently, *972SD4[1R+]* strains did not show a *SPBCPT2R1.03*. signal (also see Figure 4B).

### Cloning and sequencing of telomere-proximal *SH* regions

To overcome the difficulty of accurate assembly of repetitive sequences, ∼5 kb of the telomere-proximal part of an *SH* region in each *972SD4* where multiple common segments are aligned in a mosaic pattern (see Figure 3A), was amplified by PCR and inserted into a vector. Partial deletion series of the *SH* fragments was constructed by digesting the plasmids with restriction enzymes followed by treatment with exo- and endo-nucleases. After re-circularization of the plasmids, DNA sequences of the *SH* fragments with various lengths were determined using primers that anneal to the vector (Supplementary Figure S1, see Materials and Methods for details).

**Figure 3.**
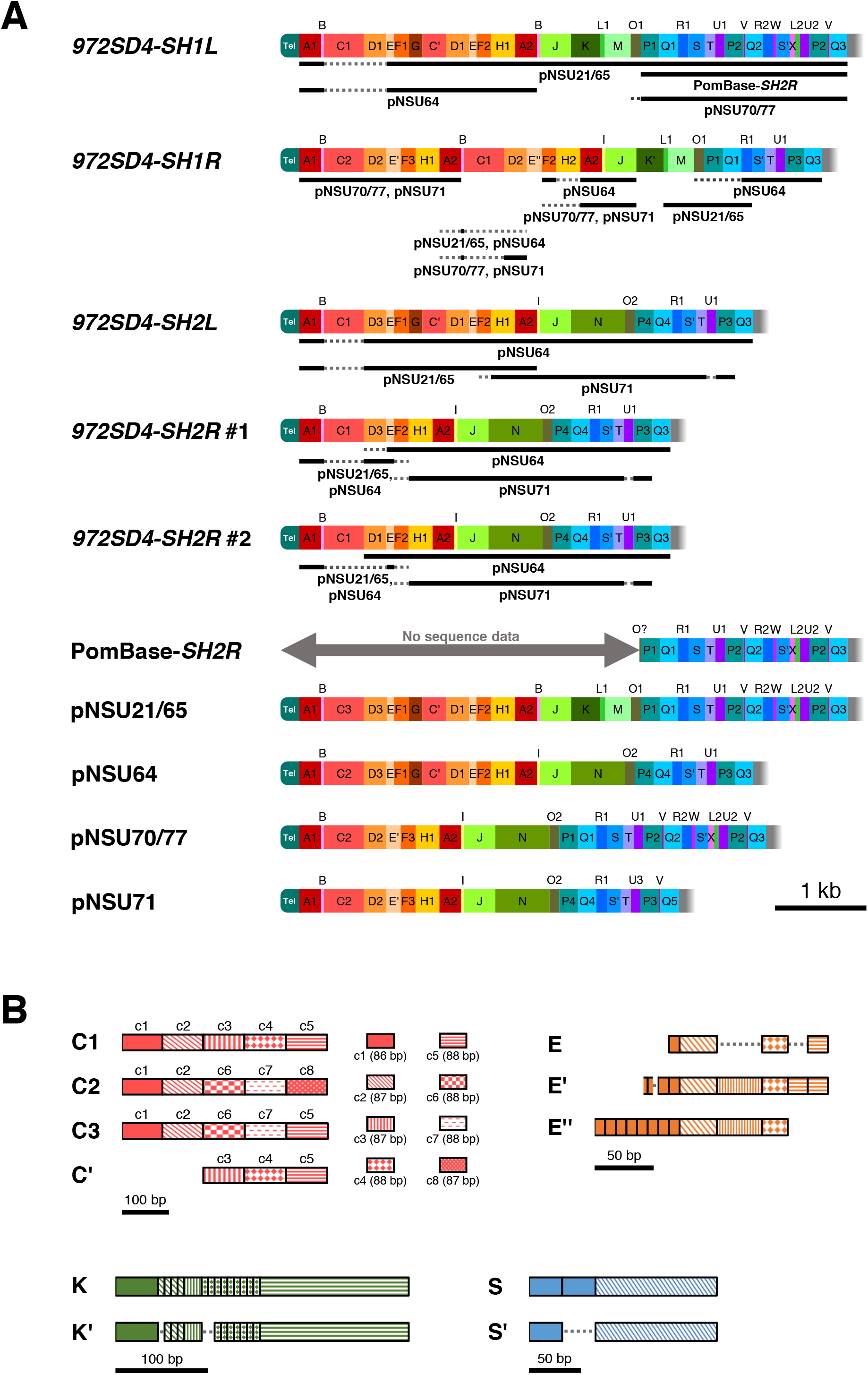
High variation in common segments of telomere-proximal *SH* regions. (A) Classification of the telomere-proximal *SH* sequences into common segments (A–X) and their variants (see Supplementary Figure S2 for each sequence). Newly sequenced *SH* regions of *972SD4* strains, a partial *SH2R* region in PomBase (PomBase-*SH2R*), and *SH* regions of a pNSU series in PomBase are shown. Note that *972SD4-SH1L*, *972SD4-SH1R*, and *972SD4-SH2L* from two independent *972SD4* strains (i.e., #1 and #2 clones) showed 100% sequence identities, whereas *972SD4-SH2R* #1 and #2 showed different variants D, D1 and D3. Black bars indicate relatively long regions that show the same segments and variants with those of pNSUs or PomBase-*SH2R*, whereas gray broken lines indicate regions that show the same segments but not the same variants. (B) Different subtypes of C, E, K, and S show different copy numbers in common sequence motifs. The same boxes show 100% sequence identity. Segment C consists of three variants, C1–3, that are composed of homologous sequence motifs, c1–8, showing >82% identities with each other (see Supplementary Figure S2C and C* for details).

### Telomere-proximal *SH* region exhibits high variation of common segments

The sequences of telomere-proximal *SH* regions of the *972SD4* strains, a part of *SH2R* in PomBase (PomBase-*SH2R*), and insertions in the pNSU series in PomBase (25) were classified into common segments (A–X) that meet the general criteria of ≥14 bp and >95% identity, with exceptional rules for subtypes C (C-1–3, see below) and C’, E, E’, and E’’, K and K’, and S and S’, which contain different copies of common sequence motifs (Figure 3B, see Materials and Methods and Supplementary Figure S2 for details). Segments were classified into variants (e.g., A1 and A2) that meet the criteria of 100% identity (Figure 3A and B, and Supplementary Figure S2). Telomere-proximal *SH* regions in two independent strains (#1 and #2) of *972SD4[1L*+], *972SD4[1R*+], and *972SD4[2L*+] exhibited 100% sequence identity, suggesting that no mutation or rearrangement had been introduced to the *SH* sequences of *972SD4* strains during amplification by PCR, cloning using *E. coli*, and serial deletion. In contrast, two strains of *972SD4[2R*+] contained different variants of segment D, D1 and D3, which showed differences in two nucleotides, suggesting that the two point mutations at segment D and/or interchromosomal recombination had occurred in *972* or *972SD4[2R*+] (Figure 3A).

We found that none of the telomere-proximal *SH* regions of *972SD4* showed the same pattern in the alignment of segments to each other; however, two pairs of regions, *972SD4-SH1L* and pNSU21/65, and *972SD4-SH2L* and pNSU64, each exhibited the same segment patterns over the whole telomere-proximal *SH* regions, suggesting the possibility that these pairs were derived from the same subtelomeres. However, the compositions of the C and D variants were different (C1 vs. C2 or C3, and D1 vs. D3). We found that segments C in *972SD4* strains were particularly different from those of pNSUs (Figure 3A, broken lines); i.e., variant C1 is the majority in the *SH* sequences of *972SD4*, whereas variant C2 is the majority in those of pNSUs. These data suggest that segment C is prone to mutations and recombination possibly due to its highly repetitive structure (Figures 3B, see Discussion).

In contrast to *972SD4-SH1L* and *972SD4-SH2L*, *972SD4-SH1R* and *972SD4-SH2R* showed combinations of pNSU patterns. Moreover, subtypes E’’ and K’, and variant H2 were unique to *972SD4-SH1R* among the *SH* sequences of *972SD4* and PomBase, suggesting that multiple times of mutation and recombination had occurred at *972SD4-SH1R*. Surprisingly, the newly sequenced *972SD4-SH2R* exhibited a pattern different from a part of the PomBase*-SH2R* sequence; rather, a part of *972SD4-SH1L* showed a pattern same as that of PomBase*-SH2R*, implying that *SH1L* and *SH2R* have exchanged their chromosomal positions over repeated rounds of cell divisions.

Although we identified several variants for each segment, these variants are not randomly combined, and partial sequences showed the same alignments; for instance, there are two common alignments for telomere-distal parts: P1-Q1-----V-Q3 (in *972SD4-SH1L*, PomBase-*SH2R*, pNSU21/65, and pNSU70/77) and E-F2-----P3-Q3 (in *972SD4-SH2L*, *972SD4-SH2R* #1, *972SD4-SH2R* #2, and pNSU64). Overall changes in the segment and variant compositions suggest that the telomere-proximal *SH* regions are hot spots for chromosome rearrangement.

### Telomere-distal *SH* regions exhibit variations with multiple insertions and deletions

We also determined sequences of telomere-distal *SH* regions in *972SD4* strains. Integration of the partial *SH* sequences in PomBase and our newly determined *SH* sequences enabled us to estimate the full length of each *SH* region: *SH1L*, 63.1 kb; *SH1R*, 40.3 kb; *SH2L*, 59.4 kb; and *SH2R*, 49.7 kb, with *SH1R* having the shortest *SH* sequence, although these lengths are likely to change through chromosome rearrangements at *SH* regions.

In contrast to the telomere-proximal *SH* regions, telomere-distal *SH* regions do not show differences in orders of homologous sequences; rather, they are highly similar (more than 99% identities with an exception at a region indicated in Figure 4A; see below). However, we found differences in length; multiple insertions or deletions were identified in these regions (Figure 4A). There are three big differences between subtelomeres (Figure 4A, thick lines). First, there is a 3.7 kb-insertion only in *SH1L* at position 4,520,104 of PomBase-*SH2R*. Second, there is a 7.1 kb-deletion only in *SH1R* at nucleotides 4,514,358–4,507,228 of PomBase-*SH2R*. This deletion was detected in the three independent strains of *972SD4[1R*+] by PFGE-Southern analysis (Figure 2D). Third, there is a 1.9 kb-deletion only in *SH1L* at nucleotides 4,500,310–4,498,437 of PomBase-*SH2R*.

**Figure 4.**
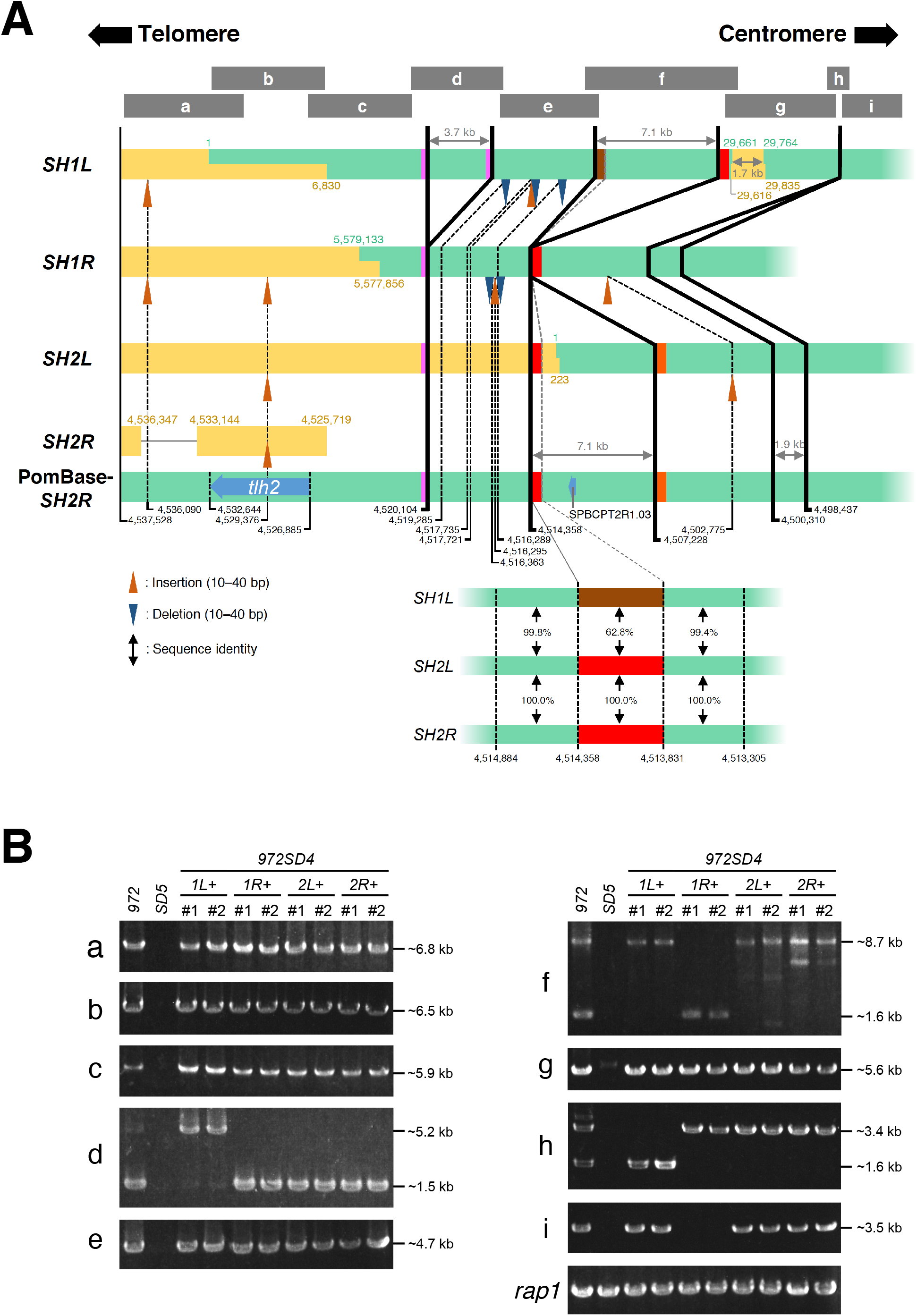
Sequence variations of telomere-distal *SH* regions. (A) Comparison of telomere-distal *SH* regions between subtelomeres. Sequences of *SH1L*, *SH1R*, and *SH2L* in PomBase (indicated by green boxes) were combined with sequences newly determined in this study (yellow boxes, showing #1 clones of each *SH* sequence), whereas newly sequenced *SH2R* fragments are shown separately with PomBase-*SH2R*. Green and yellow numbers indicate positions in each chromosome in PomBase at the ends of overlaps between PomBase and newly determined regions. The all *SH* sequences are aligned with telomeres to the left and centromeres to the right. Insertions (orange arrowheads) or deletions (blue arrowheads) of 10–40 bp are shown in comparison with PomBase*-SH2R* (thin broken lines). Insertions and deletions of <10 bp are omitted from this panel. Thick lines indicate changes in relatively long sequences. Pink, red, orange, and brown boxes indicate homologous sequence motifs at the ends of the long sequence changes (see Supplementary Figure S3 for the sequences and identities between them). Structures of *SH1L*, *SH2L*, and *SH2R* at the left end of the longest (7.1 kb) sequence change are shown below. Note that *SH1L* contains a region (a brown box) with a low identity to the other homologous sequence motifs (red and orange boxes) (see Supplementary Figure S3). Numbers in black indicate chromosomal position in PomBase-*SH2R*. Black vertical arrows indicate sequence identities. Gray boxes at the top indicate ranges of PCR products a–i analyzed in (B). (B) Lengths of the telomere-distal *SH* regions in the *972SD4* strains. DNA fragments a– i were amplified by PCR using genomic DNAs of the wild-type (JP1225), *SD5* (ST3479), and two independent *972SD4* strains (#1 and #2) as templates. Approximate DNA lengths estimated by sequences in PomBase and *972SD4* are indicated on the right.

Intriguingly, there are homologous sequences (100% identity) exist at the ends of the 3.7 kb-insertion (Figure 4A, pink boxes, and Supplementary Figure S3A), implying that these sequences had been utilized for homologous recombination (HR) when chromosomes were rearranged. Similarly, there are homologous sequences (>93% identities with one exception; see below) exist at the ends of 7.1 kb-insertions (Figure 4A, red and orange boxes, and Supplementary Figure S3A–C). However, the left-side homologous sequence of *SH1L* (*SH1L* (L) indicated by a brown box in Figure 4A) showed low sequence identity (<63%) with other homologous sequences because of multiple insertions and deletions in a region where multiple repeat sequences are arranged intricately, suggesting that chromosome rearrangements have occurred in this region (Supplementary Figure S3B and C). Interestingly, the pairs of the red boxes of *SH1L* (R) (the right-side homologous sequence of *SH1L*) and *SH1R*, the red boxes of *SH2L* (L) and *SH2R* (L), and the orange boxes of *SH2L* (R) and *SH2R* (R) show 100% sequence identity, and the difference between the pair of *SH1L* (R) and *SH1R*, and that of *SH2L* (L) and *SH2R* (L) is only one nucleotide (Supplementary Figure S3B and C). These data suggest that the 7.1 kb-deletion and/or insertion have occurred using these homologous sequences, and homologous sequence boxes (red and orange) have been copied to other *SH* regions.

There are also smaller insertions and deletions in telomere-distal *SH* regions (Figure 4A, thin broken lines for changes of 10–40 bp). Many of these are observed in no less than two *SH* regions, suggesting that these changes have been copied to other *SH* regions by chromosome rearrangement. It is noteworthy that the newly sequenced *SH2R* in *972SD4[2R*+] contains an insertion at position 4,529,376 of PomBase-*SH2R*, indicating that this insertion has been introduced to *SH2R* of strain *972* over repeated rounds of cell divisions.

### Overall structures of telomere-distal *SH* regions are stably maintained

To examine stability of the telomere-distal *SH* regions, their DNA structures in two independent strains of *972SD4[1L*+], *972SD4[1R*+], *972SD4[2L*+], and *972SD4[2R*+] were analyzed by PCR using multiple primer sets (PCR products a–i are indicated by gray bars in Figure 4A, top). We found that lengths of the all PCR products matched those speculated by PomBase (Figure 4B), indicating that the overall DNA structures of telomere-distal *SH* regions have been relatively stably maintained compared with those of telomere-proximal *SH* regions after repeated rounds of cell division.

### *SH* regions are hot spots for mutations

Given that the newly sequenced telomere-proximal *SH* regions of *972SD4* contain multiple nucleotide changes compared with those in PomBase (Figure 3), we examined whether mutation rates are specifically high in *SH* regions. Sequences of multiple Ch2 loci in two independent *972SD4[2R*+] strains were determined and compared with those of PomBase (Figure 5 and Supplementary Table S3). We found that *SH* regions, especially around the *tlh2* gene locus, exhibit high rates of mutations. Most of them are point mutations, but some are changes of numbers of repeat sequences such as [T]n and [ACAACG]n. In contrast, the telomere-distal half of *SH* region and all chromosomal regions outside of *SH*, i.e., the *SH*-adjacent region within the subtelomere (the knob region), the subtelomere boundary region, the subtelomere-adjacent region, and the *rap1* locus, which is far from the subtelomere, show 100% identities with sequences in PomBase, indicating strict preservation of their DNA sequences through repeated rounds of cell division. These data suggest that the telomere-proximal half of *SH* regions (∼25 kb) are particularly prone to the accumulation of mutations.

**Figure 5.**
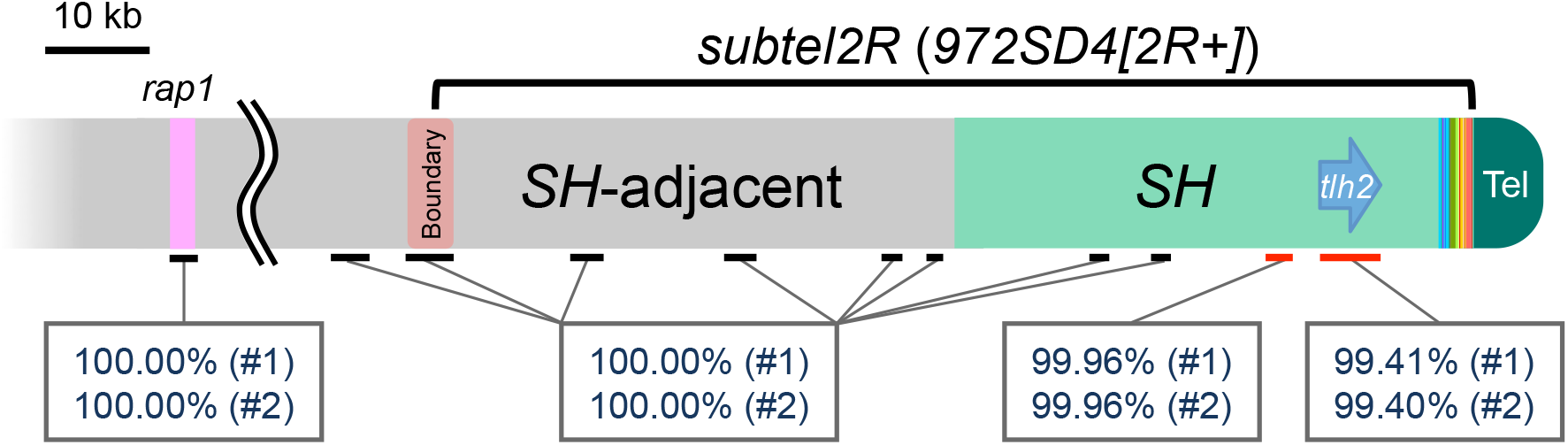
Sequence changes in the telomere-distal *SH2R* region of *972SD4[2R*+] compared with that in PomBase. Summary of sequence identities between *2R* (the right arm of Ch2) of two independent *972SD4[2R+]* strains (clones #1 and #2) and that of PomBase. Sequences of the telomere-distal *SH2R* region, the *SH2R-*adjacent subtelomere region (the knob region), the subtelomere boundary, the subtelomere-adjacent euchromatin region, and the *rap1* locus were analyzed. The *rap1* locus is located ∼1.4 Mb from the telomere of *2R*. Red bars indicate regions with nucleotide changes compared with those in PomBase (see Supplementary Table S3 for details).

### Identification of new members of the subtelomeric RecQ-type DNA helicase gene family

In genome sequences in PomBase, there are two RecQ-type DNA helicase genes, *tlh1* (partial) and *tlh2*, which have been allocated to *SH1L* and *SH2R*, respectively. Parts of the DNA sequences of *tlh1/2* are homologous with the *dh* repeat sequence of pericentromeres and serve as templates for small interfering RNA (siRNA) produced by RNA interference (RNAi) machinery; further, the siRNA participates in the initiation of subtelomeric heterochromatin formation (22,23). Our sequencing data of *SH* regions newly identified two members of the *tlh* gene family, *tlh3* and *tlh4*, in *SH1R* and *SH2L*, respectively. Thus, *S. pombe* genome contains four *tlh* genes in total (Figure 6A).

**Figure 6.**
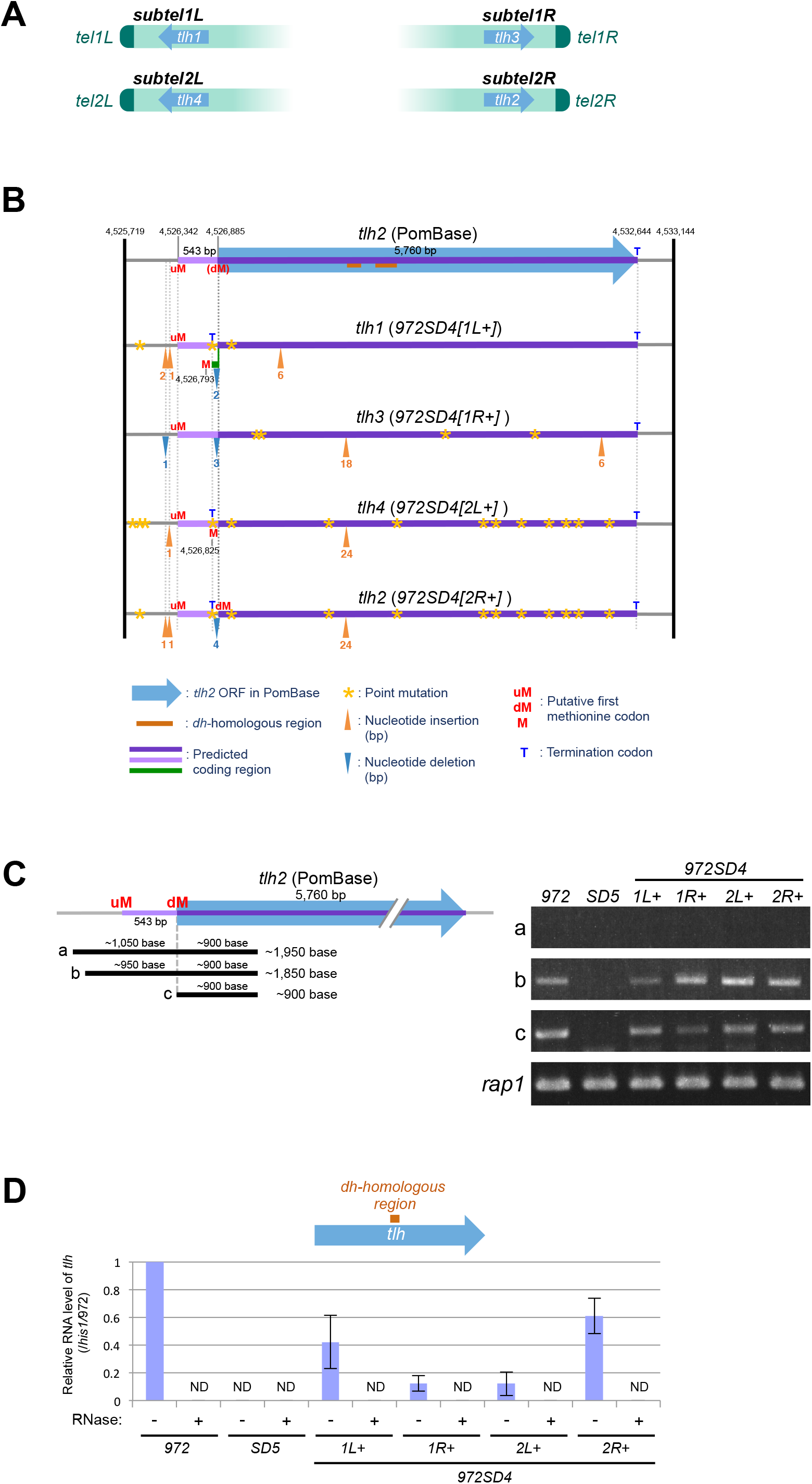
Identification of new *tlh* genes with multiple mutations. (A) Identification of *tlh3* and *tlh4*. Locations of the four *tlh* genes on chromosomes are shown. (B) Summary of predicted ORFs and nucleotide changes in the *tlh* genes of *972SD4* compared with the *tlh2* gene in PomBase. The top panel shows the *tlh2* locus in PomBase. Seven-digit numbers indicate chromosomal positions on Ch2 in PomBase. Blue arrow with a purple line, the *tlh2* ORF defined in PomBase; brown bar, *dh*-homologous region; pale purple line, predicted additional coding region in the reading frame same as that of the *tlh2* ORF in PomBase; green line, predicted additional coding region in the reading frame different from that of the predicted additional *tlh2* ORF in PomBase (a pale purple line); yellow star, point mutation (no frame shift); orange arrowhead, nucleotide insertion (number of inserted nucleotides is shown below); blue arrowhead, nucleotide deletion (number of deleted nucleotides is shown below); dM, the original first methionine codon of the *tlh2* ORF in PomBase; uM, putative first methionine codon upstream of dM; M, putative first methionine codon between uM and dM; T, terminal codon. (C) Expression of RNA from various regions (a–c) of the *tlh* locus and the *rap1* locus (control) in strains *972*, *SD5*, and *972SD4* was analyzed by PCR. The PCR products were analyzed by gel electrophoresis. (D) Expression of RNA from the *dh*-homologous region of the *tlh* genes in strains *972*, *SD5*, and *972SD4* was analyzed by quantitative RT-PCR in the presence (+) or absence (–) of RNase. Expression of *tlh*^+^ relative to that of *his1*^+^ was normalized to that in *972* (–RNase). Error bars indicate the standard deviations. ND, not detected. *N* = 3.

### The *tlh* genes in *972SD4* contain multiple nucleotide changes

Examination of the *tlh* sequences raised the possibility that lengths of their ORFs are different from that defined for *tlh2* in PomBase. In the sequence of PomBase-*SH2R*, there is another methionine (Met) codon (uMet [<u>u</u>pstream <u>Met</u>]) 543 bases (corresponding to 181 amino acids) upstream from the original first Met codon (dMet [<u>d</u>ownstream <u>Met</u>]) that was defined by PomBase in the same reading frame (Figure 6B). Moreover, the sequences containing these Met codons match the consensus Kozak sequence (A/GNN<u>ATG</u>G, the first methionine codon underlined), which participates in the initiation of translation in eukaryotes (32).

Unexpectedly, each of the newly sequenced *tlh1–4* genes of *972SD4* was found to contain multiple nucleotide changes (i.e., point mutations, insertions, and deletions) compared with the *tlh2* gene in PomBase (Figure 6B). There are tandem repeat sequences at most of the loci where insertion or deletion occurred (Figure 6B, orange and blue arrowheads), suggesting that these repetitive sequences are highly prone to recombination. Because of these changes, three of the genes (*tlh1*, *tlh2*, and *tlh4*) have a putative short coding sequence (480 bases) between the uMet codon and a termination codon derived by a point mutation (Figure 6B, pale purple lines), in addition to the long coding sequence homologous to that of *tlh2* in PomBase (Figure 6B, purple lines with or without a green line). Intriguingly, multiple nucleotide changes, including point mutations, insertions, and deletions, did not introduce premature termination codons within the long ORFs of the *tlh* genes in *972SD4*.

### RNA expression from the *tlh* genes in *972SD4*

To examine our assumption that the *tlh* genes have ORFs longer than that previously predicted by PomBase, we determined ranges of *tlh* RNA expression by RT-PCR (Figure 6C). We found that RNAs of *tlh* genes were expressed from at least 950 bases upstream of the dMet codon (∼400 bp upstream of the uMet codon). These data suggest that the *tlh* genes may have ORFs that are potentially 543 bases longer than that defined in PomBase (Figure 6B, a pale purple line in the top panel).

To examine whether the *tlh* genes in *972SD4* carrying multiple mutations are capable to produce *dh* RNAs to participate in the formation of subtelomeric heterochromatin, we determined RNA expression from the *dh*-homologous region of each *tlh* gene by RT-PCR. We found that these four *tlh* genes express *dh* RNAs; however, the expression levels were lower than that in the wild-type strain and variable between the *tlh* genes probably because of unstable epigenetic gene silencing by the formation of subtelomeric heterochromatin and multiple mutations (Figure 6D). These results suggest that DNAs of the *tlh* genes are prone to mutations without affecting RNA expression.

## Discussion

In this study, we obtained complete sequences of *S. pombe* subtelomeres by producing strains with single *SH* regions. The whole sequences revealed that *SH* regions are composed of two parts: the ∼5 kb telomere-proximal region and ∼35–57 kb telomere-distal region. The telomere-proximal region is a mosaic of multiple common segments that vary between subtelomeres and strains, suggesting that this region is highly prone to chromosome rearrangement during cell divisions. In contrast, the telomere-distal region shows high sequence similarity between subtelomeres and strains, although there are some insertions and deletions, suggesting that the overall DNA structure of this region is maintained stably. Intriguingly, the telomere-proximal half of *SH* region exhibits relatively high rates of nucleotide mutations, whereas chromosomal regions outside of this region show strict genome preservation. Thus, *SH* regions exhibit multiple patterns of genome variation (Figure 7).

**Figure 7.**
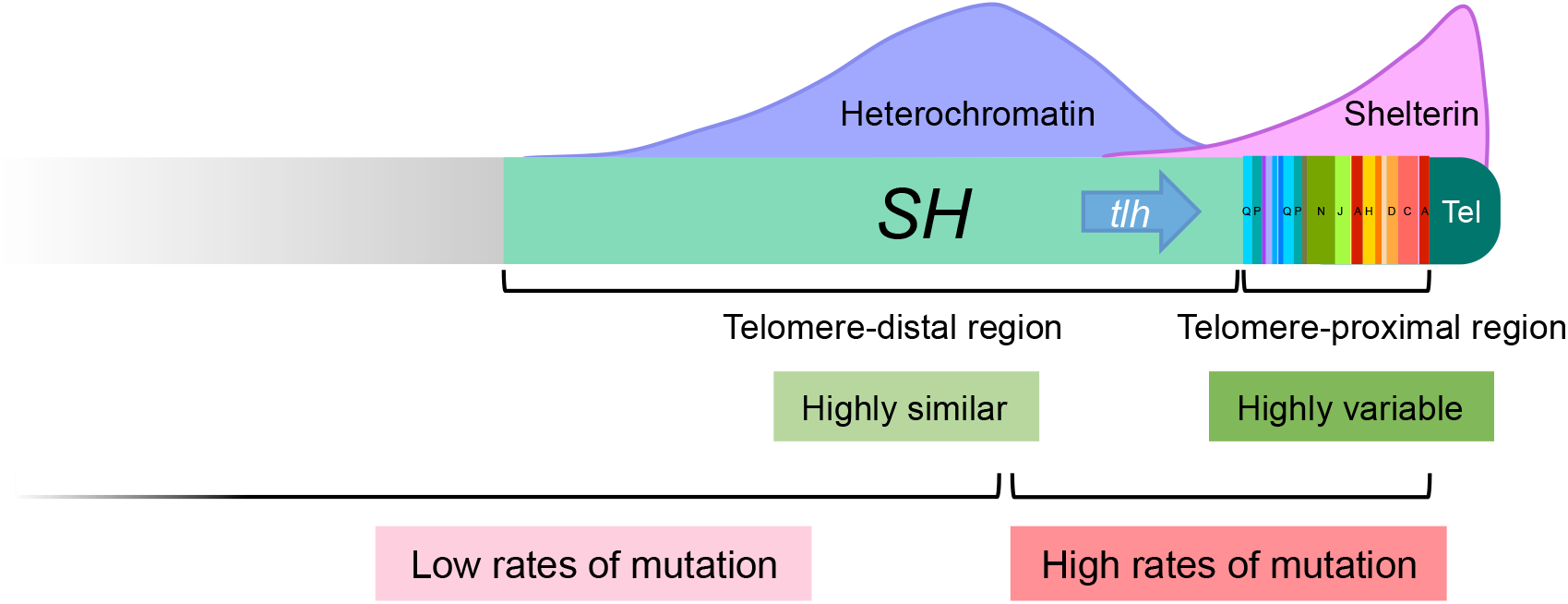
Summary of structures and features of the *SH* region in *S. pombe*. The *S. pombe SH* region is composed of two parts. The telomere-proximal *SH* region is a mosaic of multiple common segments that vary between subtelomeres and strains. On the other hand, the telomere-distal *SH* region contains sequences highly similar between subtelomeres despite containing deletions and insertions. The telomere-proximal half of *SH* region (∼25 kb) show high rates of mutation, whereas the telomere-distal half of *SH* region (∼25 kb) and chromosomal loci that are outside of *SH* show very low mutation rates. Previous studies showed that the telomere protein complex called shelterin associates with telomeres and spread to the telomere-proximal *SH* region (34). The formation of subtelomeric heterochromatin is initiated by shelterin and RNAi, the latter of which utilizes RNA molecules from the *tlh* genes, and it spread to *SH* regions (21,23).

Why do telomere-proximal and telomere-distal *SH* regions exhibit these distinct features? We previously reported that the telomere protein complex called shelterin, which plays crucial roles at telomeres (15,33), is localized at the telomere-proximal *SH* regions, in addition to telomeres (23,34), whereas subtelomeric heterochromatin is localized mainly at the telomere-distal *SH* regions (21) (Figure 7). Heterochromatin structures generally repress recombination (35); therefore, it is supposed that repression of recombination by subtelomeric heterochromatin brings about stability of overall structures of the telomere-distal *SH* regions. In contrast, it is likely that shelterin allows recombination, although telomeres and telomere-proximal *SH* regions form compact structures showing strong gene silencing (23,36,37).

Human subtelomeres (*SH* regions in humans) are also mosaics of multiple common segments corresponding to the telomere-proximal region of *S. pombe*, but they contain no sequence equivalent to that of the telomere-distal *SH* region, i.e., a relatively long common sequence shared by all subtelomeres (9,19). Common segments of the same categories are mostly non-identical (∼90–100% identities), and the location and copy number of each segment vary among individuals (9). In *S. cerevisiae*, the subtelomeres show common X and Y’ elements, and ORFs of proteins such as PAU and FLO families; however, copy numbers of the Y’ element and the ORFs are highly variable between strains (18,38). Based on these findings and studies in other species, along with our results in *S. pombe*, we propose that high variation in *SH* sequences are common phenomenon in eukaryotes.

What underlies this high variation of subtelomeres? First, DNA double-strand breaks (DSBs) are repaired by either HR or non-homologous end joining (NHEJ). Vegetative cell cycle of the wild-type *S. pombe* strain lacks G1 phase because cells after mitosis possess sufficient mass to proceed to S phase. Therefore, ∼80% period of the *S. pombe* cell cycle is G_2_ phase, when HR predominates for DSB repair (39). Because of the high identities between *SH* regions, DNA repair by HR may occur frequently between different chromosomes (interchromosomal repairs), as well as between sister chromosomes (intrachromosomal repairs), causing gross rearrangement of chromosomes. Second, repetitive sequences within *SH* regions may be recognized by HR machineries, causing amplification or deletion of repeat units. Third, repetitive sequences, including *S. pombe* telomeres and subtelomeres, are regions intrinsically difficult to replicate during S phase (40,41). Replication fork collapse and erosion of telomeres and subtelomeres can result in formation of one-ended DNA breaks that are repaired by break-induced replication (BIR) (42). Recent studies suggested that BIR is highly inaccurate repair and causes high levels of mutations and chromosome rearrangements (43–48). Therefore, BIR may cause high rates of mutations and chromosome rearrangements in *SH* regions.

Taken together, we propose that subtelomeres are highly polymorphic chromosomal regions and contribute to genome evolution. Importantly, in the *SH* regions of *S. pombe*, there is no gene essential for cell growth under normal culture conditions, and deletion of all of *SH* regions does not affect cell growth *per se* (21). Genomes of other species also contain multiple copies of the same genes at *SH* regions. These are why cells can continue to grow, even with mutations in *SH* regions, resulting in the accumulation of mutations. In this study, we have shown that the sequence of the *tlh2* gene has changed from that of the original strain *972* after repeated rounds of cell division (Figure 6B). Human *SH* regions contain various genes (9,19). Thus, the high rates of polymorphisms in *SH* regions may contribute to human diversity and sometimes to disease susceptibilities. Overall, genome rearrangement, deletion, insertion, and mutation can cause changes in ORFs, which will result in diversification of species.

## Data availability

DNA sequences of newly sequenced *SH* regions are available in the DNA Data Bank of Japan (DDBJ) under accession codes LC521649 (∼1.7 kb of the telomere-distal *SH1L* in *972SD4[1L*+] #1), LC521650 (∼1.7 kb of the telomere-distal *SH1L* in *972SD4[1L*+] #2), LC521651 (the telomere-proximal *SH1L* in *972SD4[1L*+] #1), LC521652 (the telomere-proximal *SH1L* in *972SD4[1L*+] #2), LC521653 (the telomere-proximal *SH1R* in *972SD4[1R*+] #1), LC521654 (the telomere-proximal *SH1R* in *972SD4[1R*+] #2), LC521655 (the telomere-proximal *SH2L* in *972SD4[2L*+] #1), LC521656 (the telomere-proximal *SH2L* in *972SD4[2L*+] #2), LC521657 (the telomere-proximal *SH2R* in *972SD4[2R*+] #1), and LC521658 (the telomere-proximal *SH2R* in *972SD4[2R*+] #2).

## Acknowledgements

We thank H. Maekawa for critical reading of the manuscript, T. Nakagawa for valuable comments, and all of the lab members for discussion and support.

## Author contributions

J.K. conceived the project, T.K., S.T., and J.K. designed and performed experiments. Y.O., T.K., and J.K. analyzed data. J.K., Y.O., and T.K. wrote the manuscript. Y.T. contributed to sequencing of *SH* regions.

## Supplementary information

### Supplementary Figures

**Figure S1.**
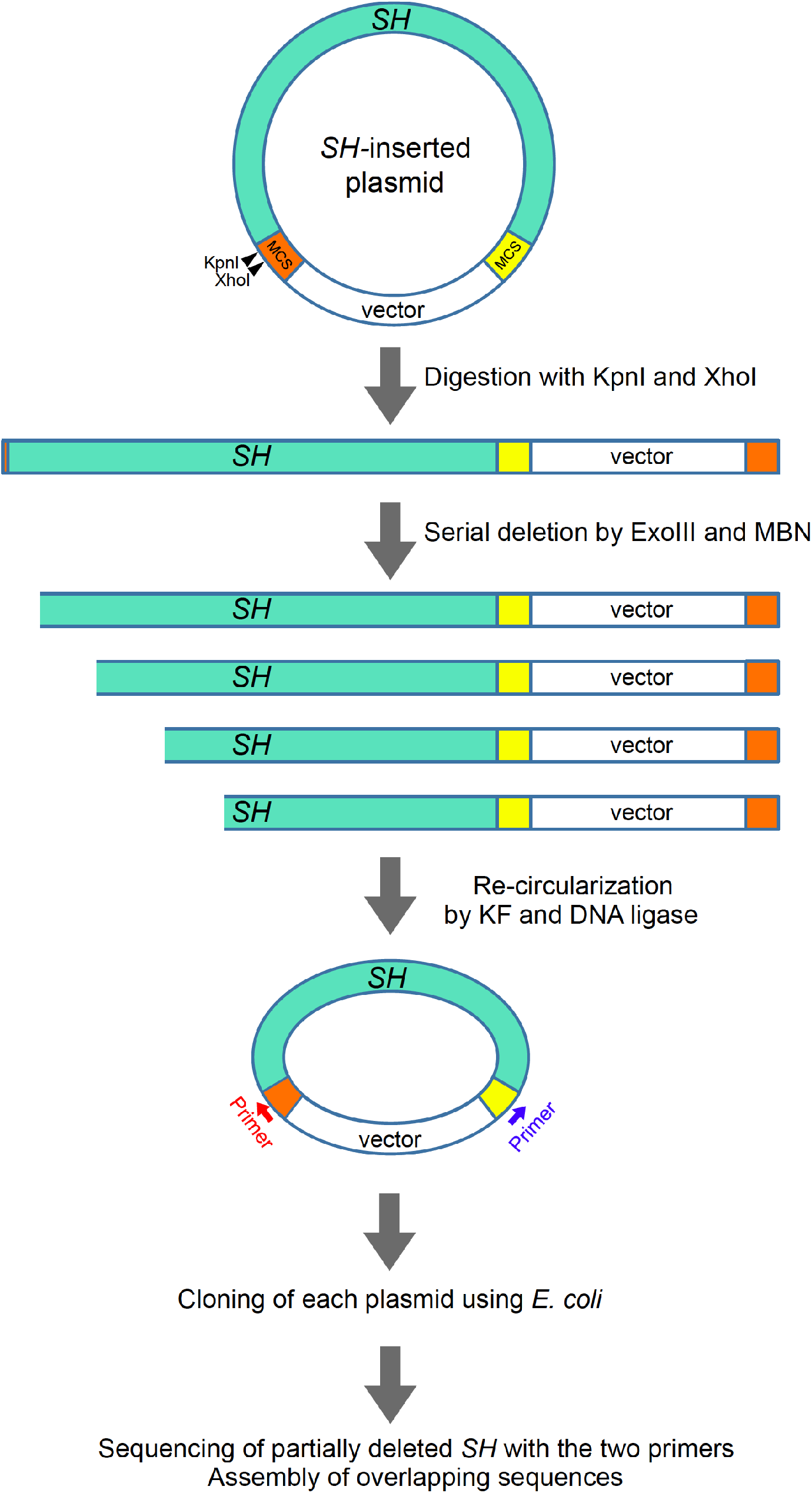
Sequencing of telomere-proximal *SH* regions by the serial deletion method. A telomere-proximal *SH* region with various common segments (∼5 kb) was amplified by PCR from each *972SD4* strain, and inserted into a vector. The resultant plasmid was digested with restriction enzymes, KpnI and XhoI, at the multi-cloning sites (MCSs) of the vector. The linearized plasmid was serially deleted from the 5’-protruding XhoI cutting site by treatment with exonuclease III (ExoIII) and mung bean nuclease (MBN) for fixed times. After deletion of the *SH* region, both ends of the plasmid were blunted by klenow fragment (KF) and ligated by DNA ligase. The resultant re-cirsularized plasmids were cloned using *E. coli*. The partially deleted *SH* region was sequenced using primers that aneal to MCSs of the vector. And the sequences were assembled using overlapping sequences.

**Figure S2.**
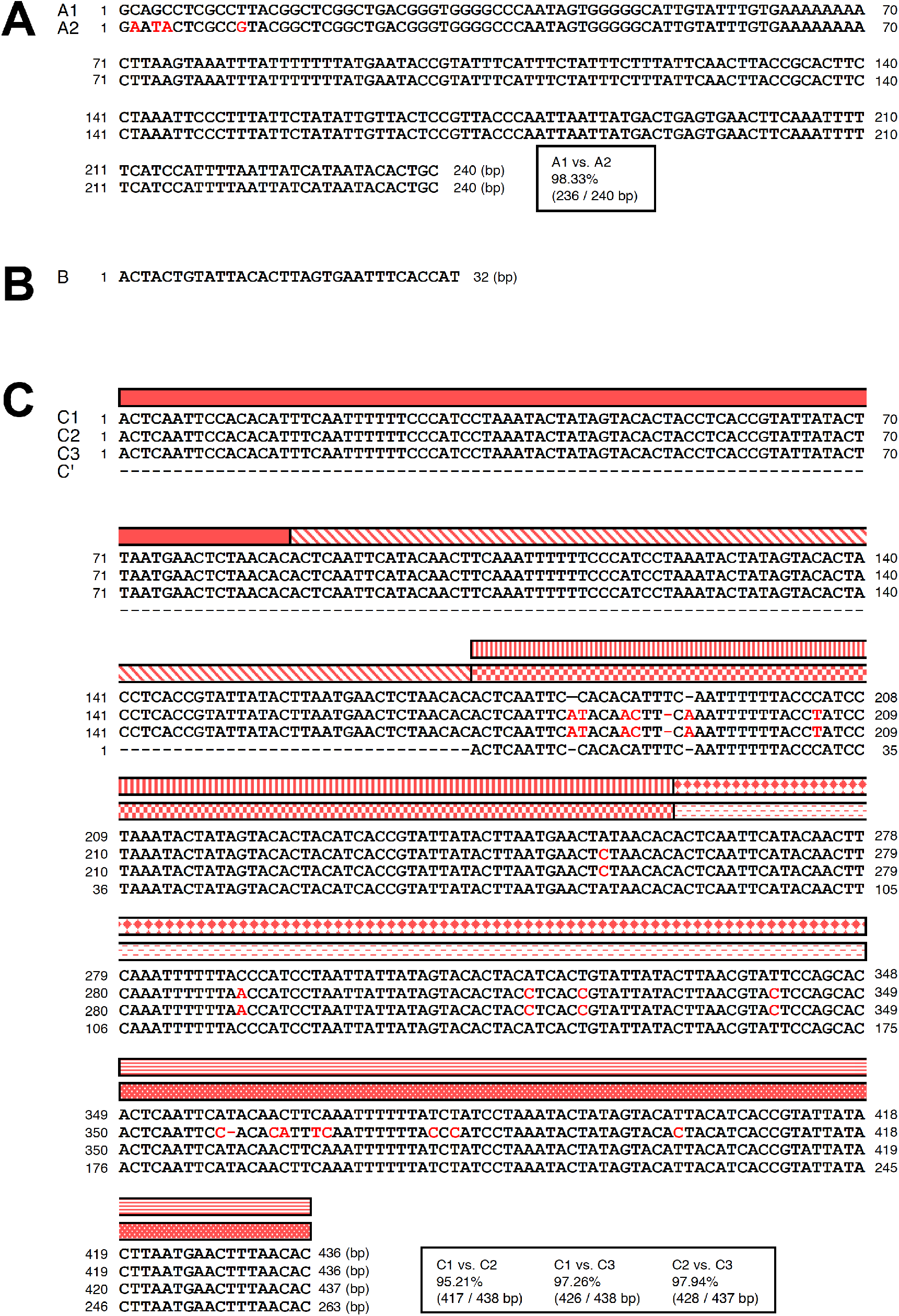
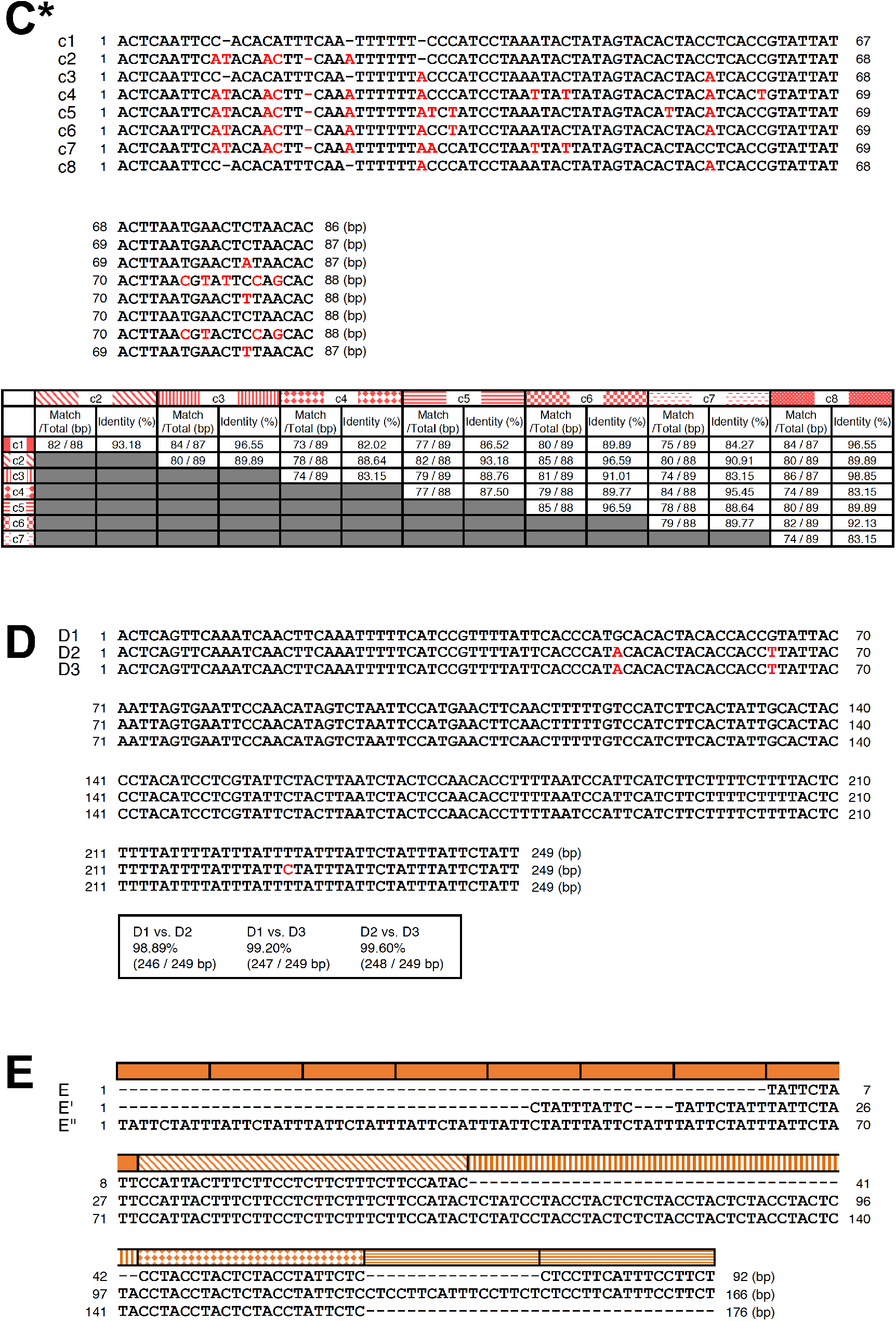
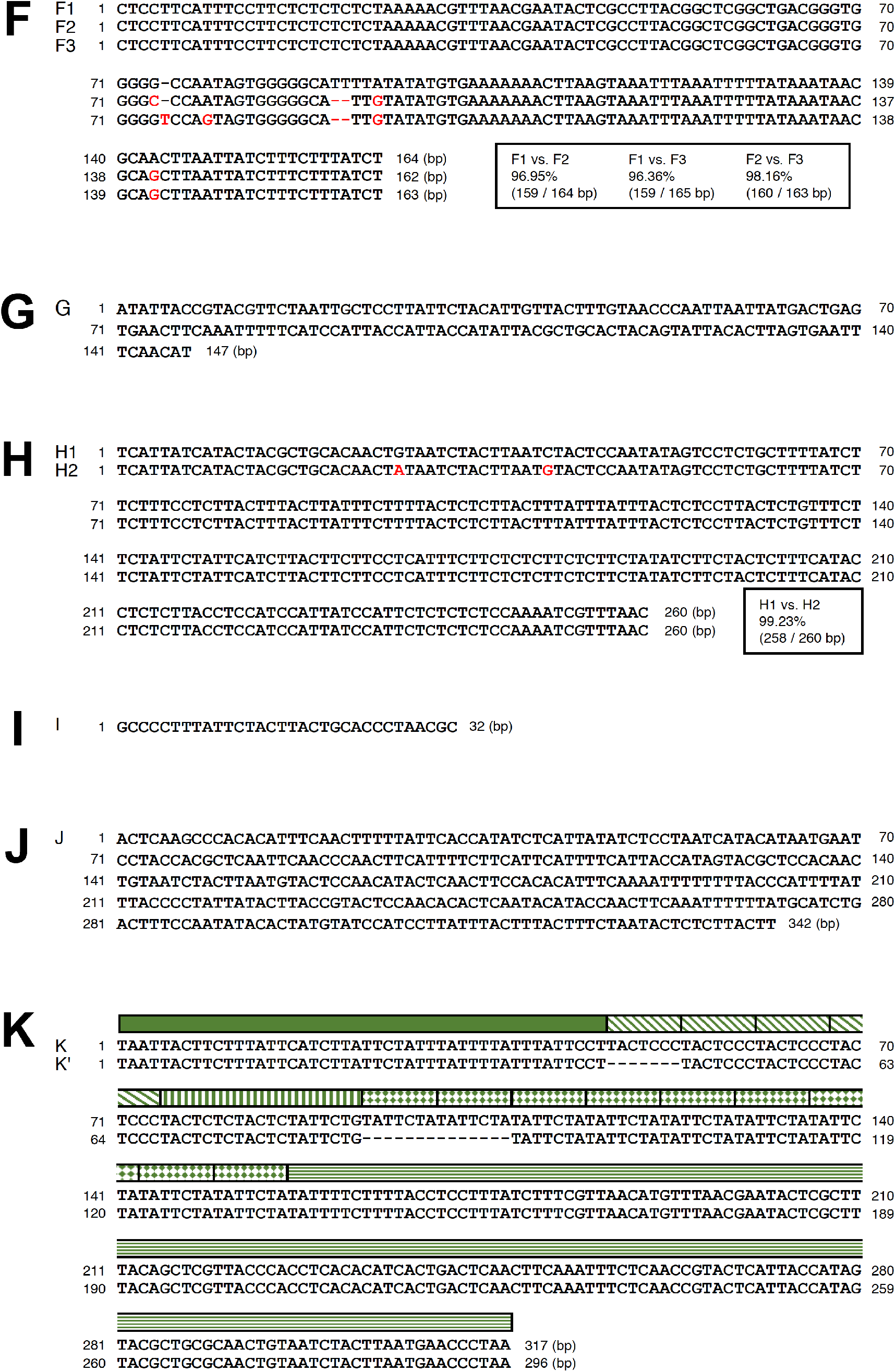
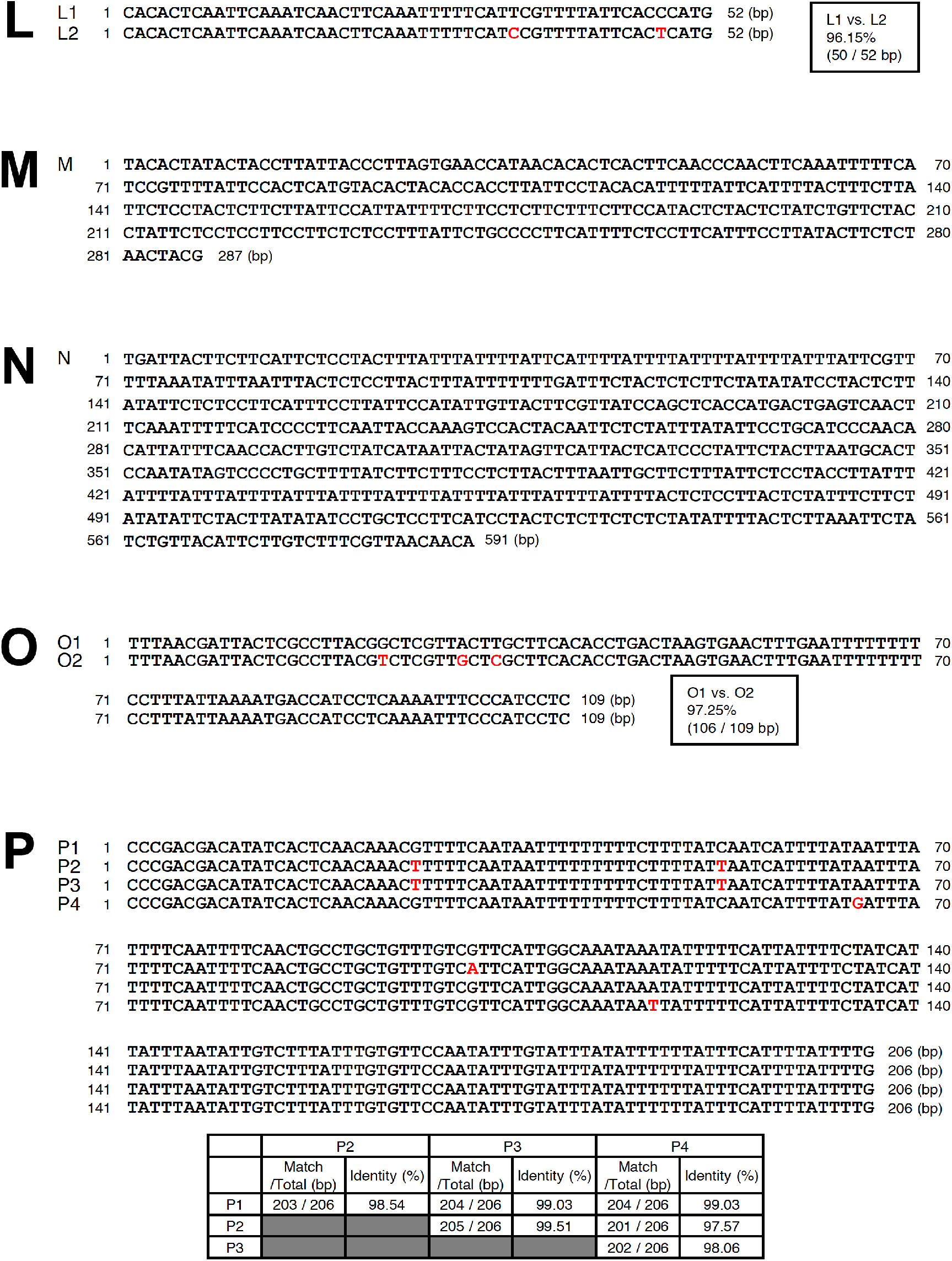
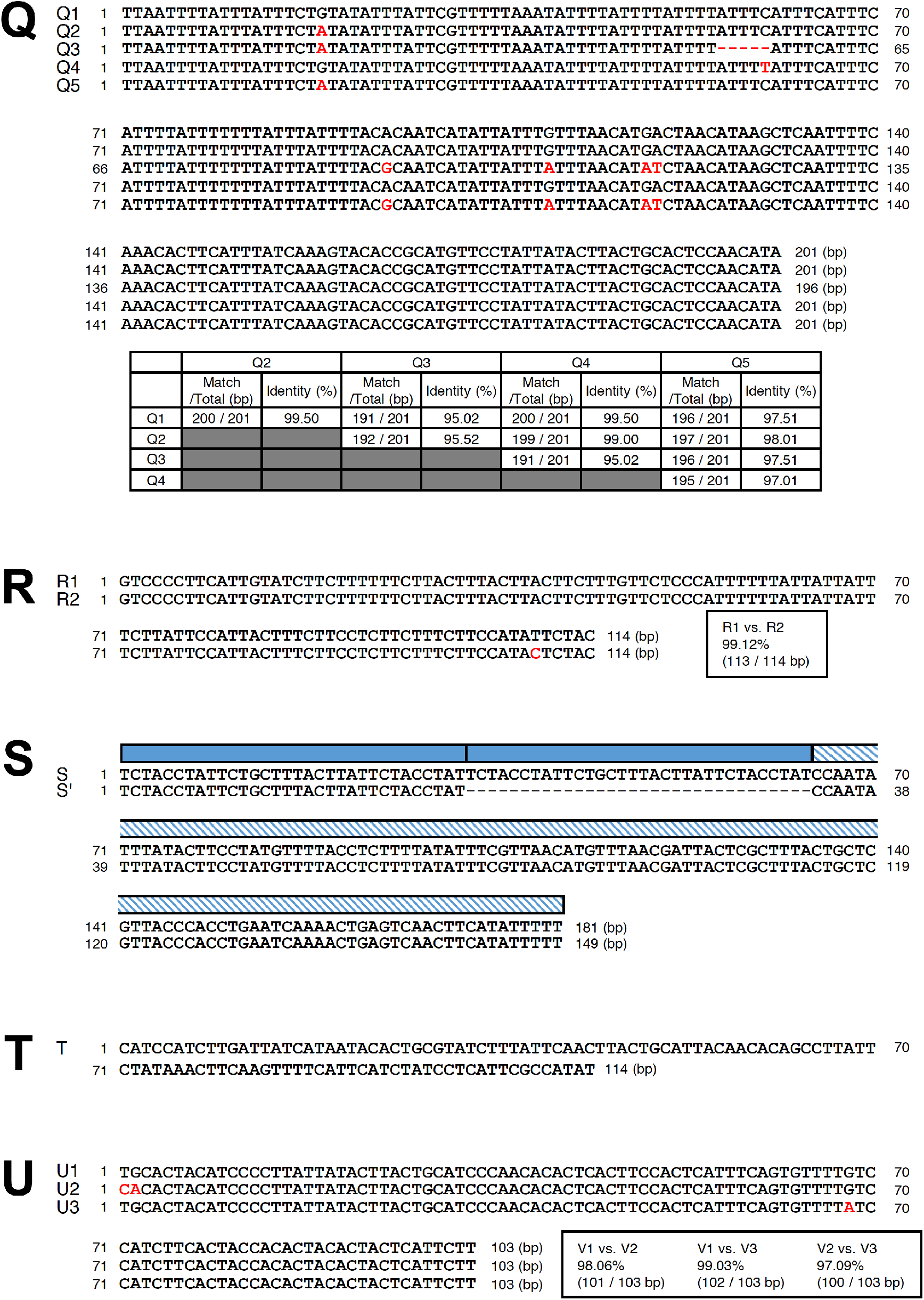
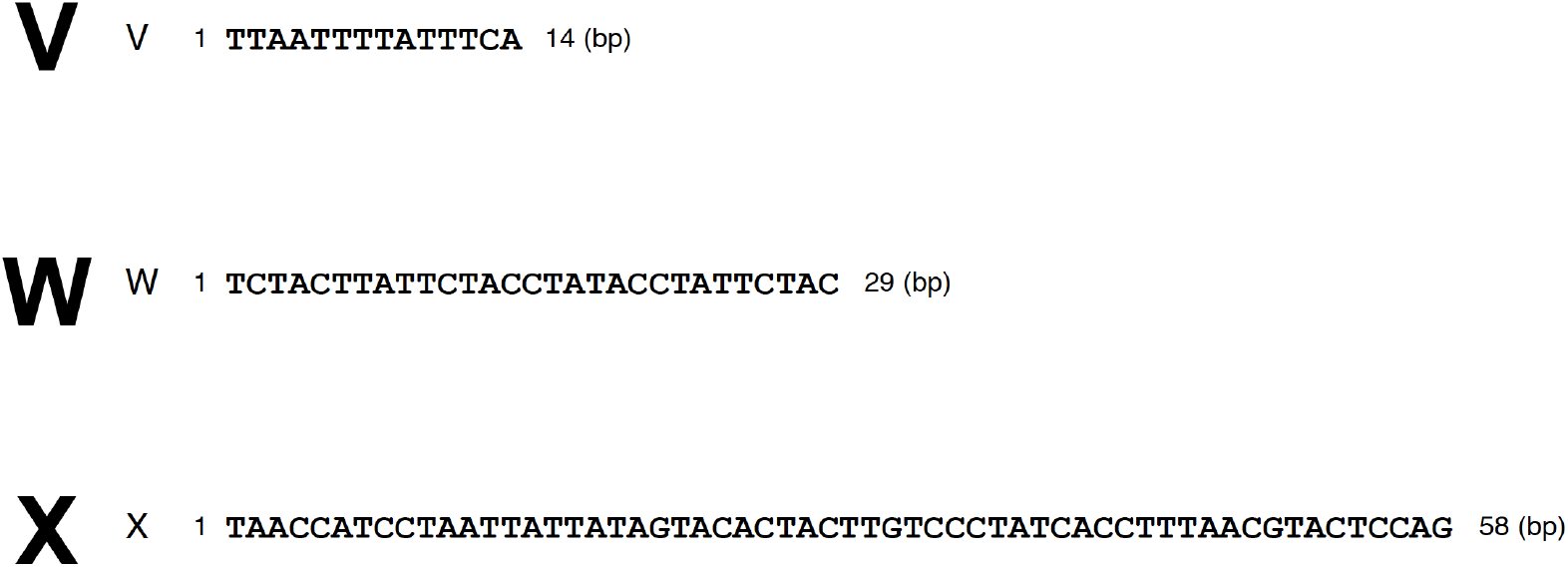
Sequences of common segments in telomere-proximal *SH* regions. Sequences of subtypes and variants of each segment are aligned. Nucleotide changes are shown in red. Sequence identities between variants of each segment are indicated. Positions of common sequence motifs in segments C, E, K, and S are indicated according to Figure 3B. Panel C* shows sequences of the common sequence motifs of segment C, c1–8, and identities between them.

**Figure S3.**
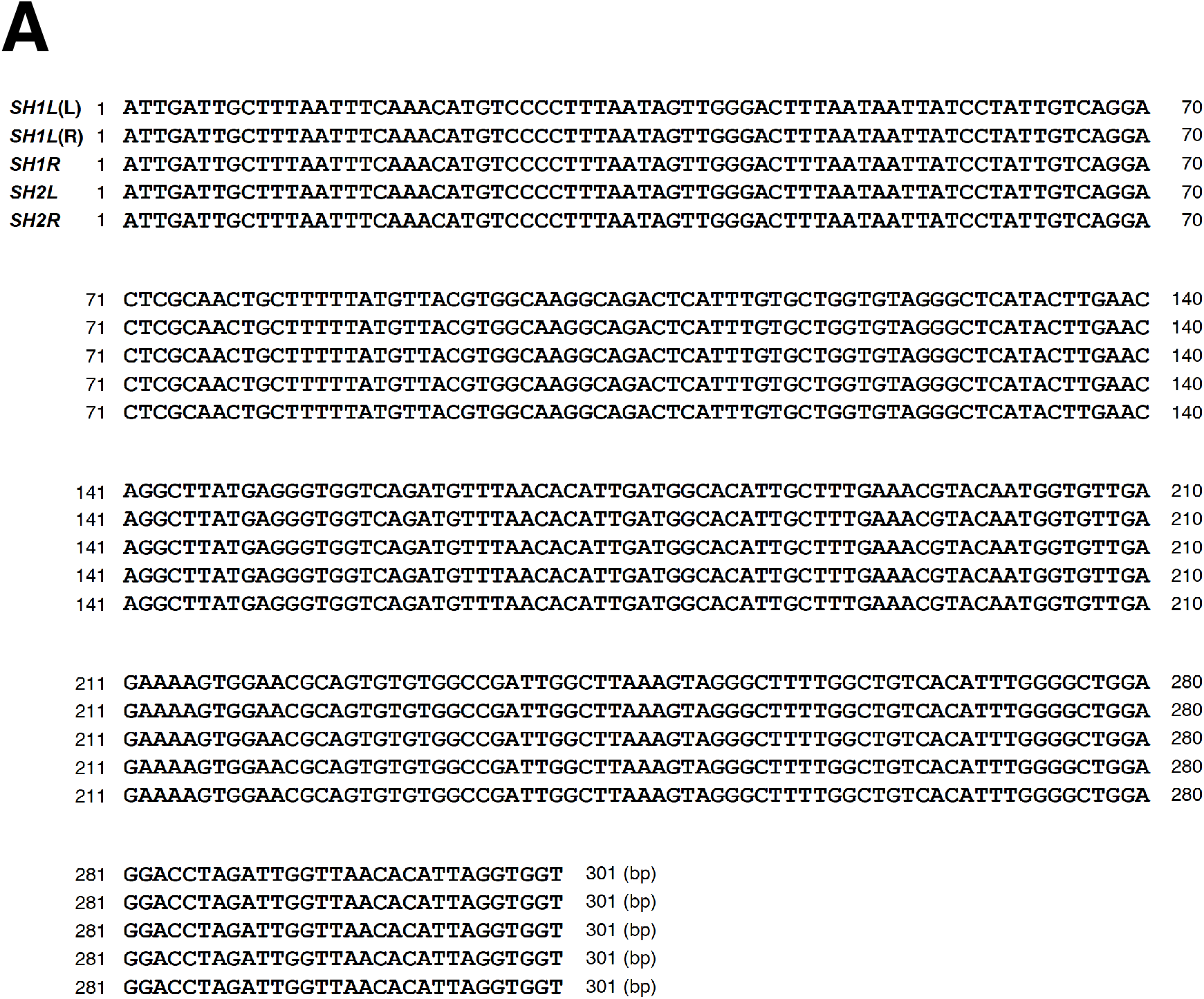
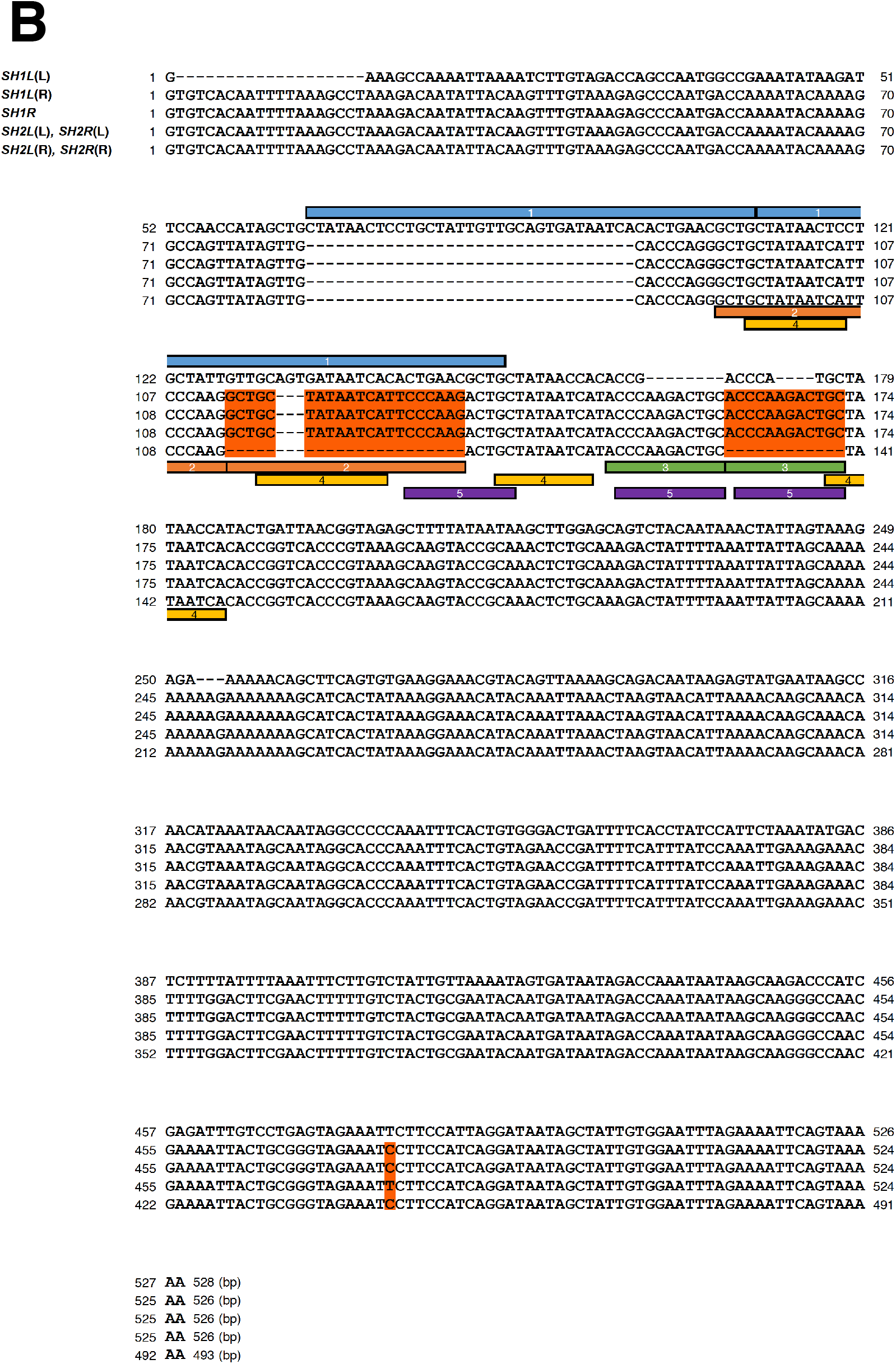
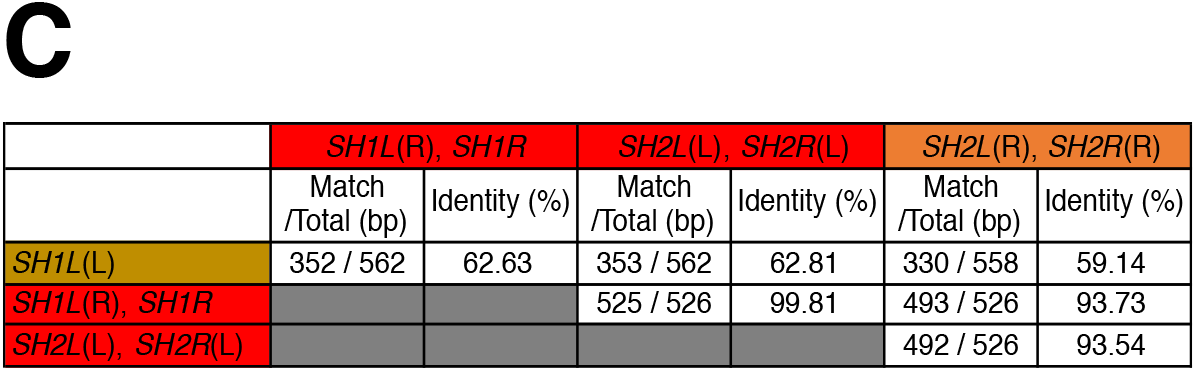
Homologous sequences at the ends of insertions in telomere-distal *SH* regions. (A) Homologous sequences at the ends of the 3.7 kb-insertion (pink boxes in Figure 4A). *SH1L* (L), a homologous sequence on the left side of the 3.7 kb-insertion in *SH1L* in Figure 4A; *SH1L* (R), that on the right side of the 3.7 kb-insertion in *SH1L*; *SH1R*, that in *SH1R*; *SH2L*, that in *SH2L*; *SH2R*, that in *SH2R*. Note that these sequences are 100% identical. (B) Homologous sequences at the ends of the 7.1 kb-insertion (red, orange, and brown boxes in Figure 4A). Note that the region containing multiple repeat units (1–5) shows high sequence variation. Sequence variations among *SH1L* (R), *SH1R, SH2L* (L), *SH2R* (L), *SH2L* (R), and *SH2R* (R) are highlighted in orange. (C) Sequence identities between homologous sequences in (B). Note that sequences of the pairs, *SH1L* (R) and *SH1R, SH2L* (L) and *SH2R* (L), and *SH2L* (R) and *SH2R* (R) are 100% identical.

### Supplementary Tables

**Table S1.**
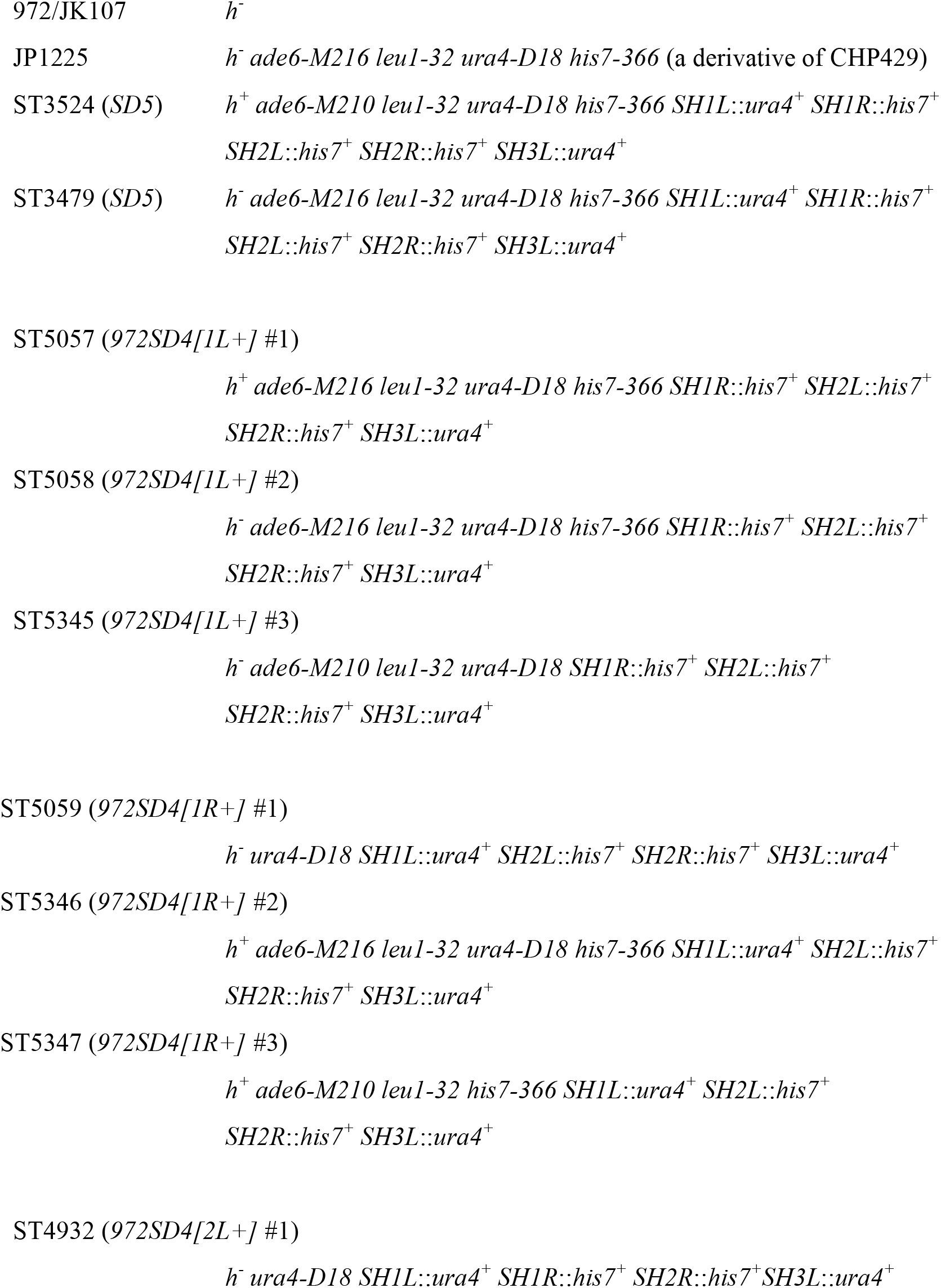
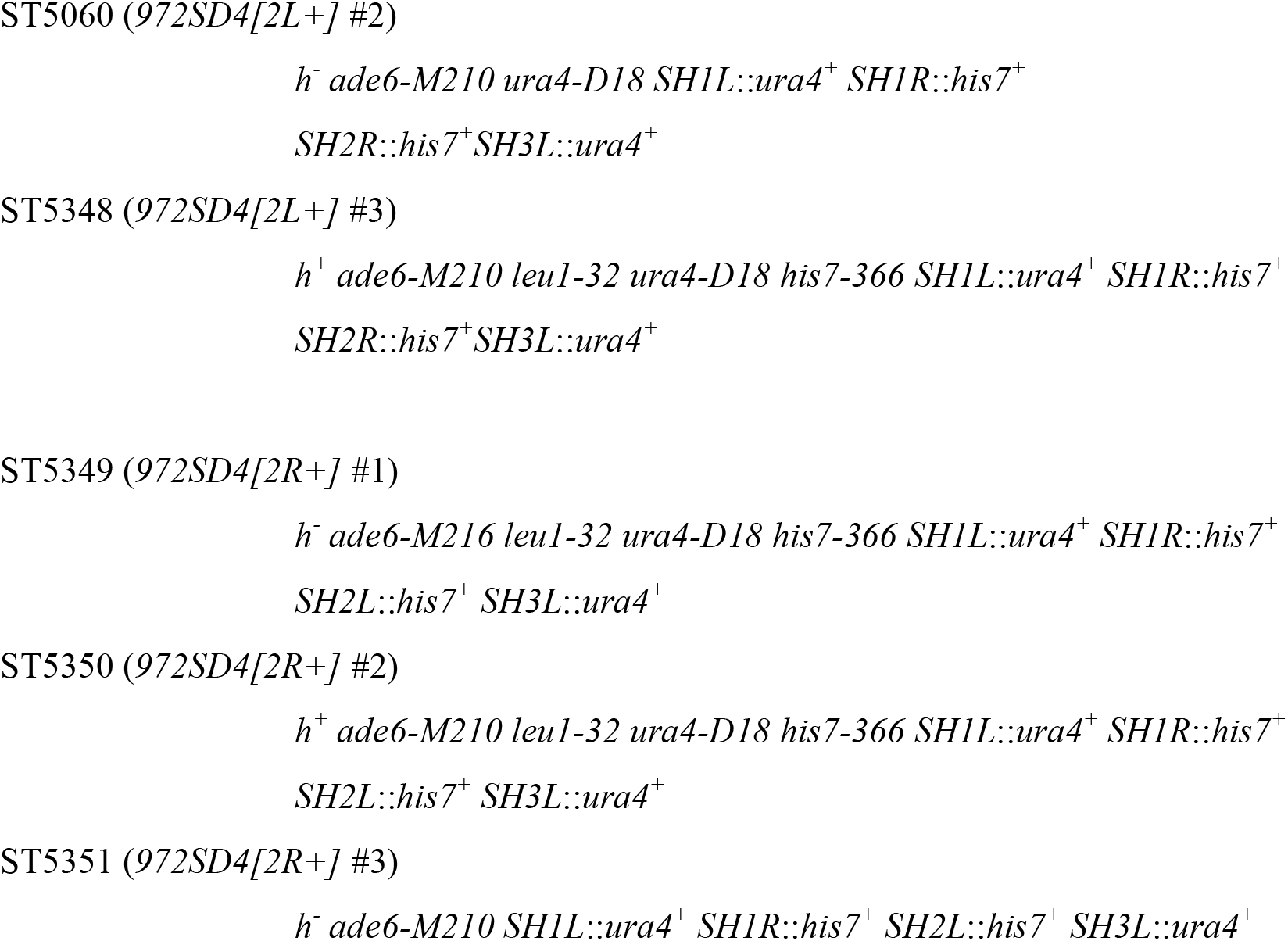
Fission yeast strains used in this study.

**Table S2.**
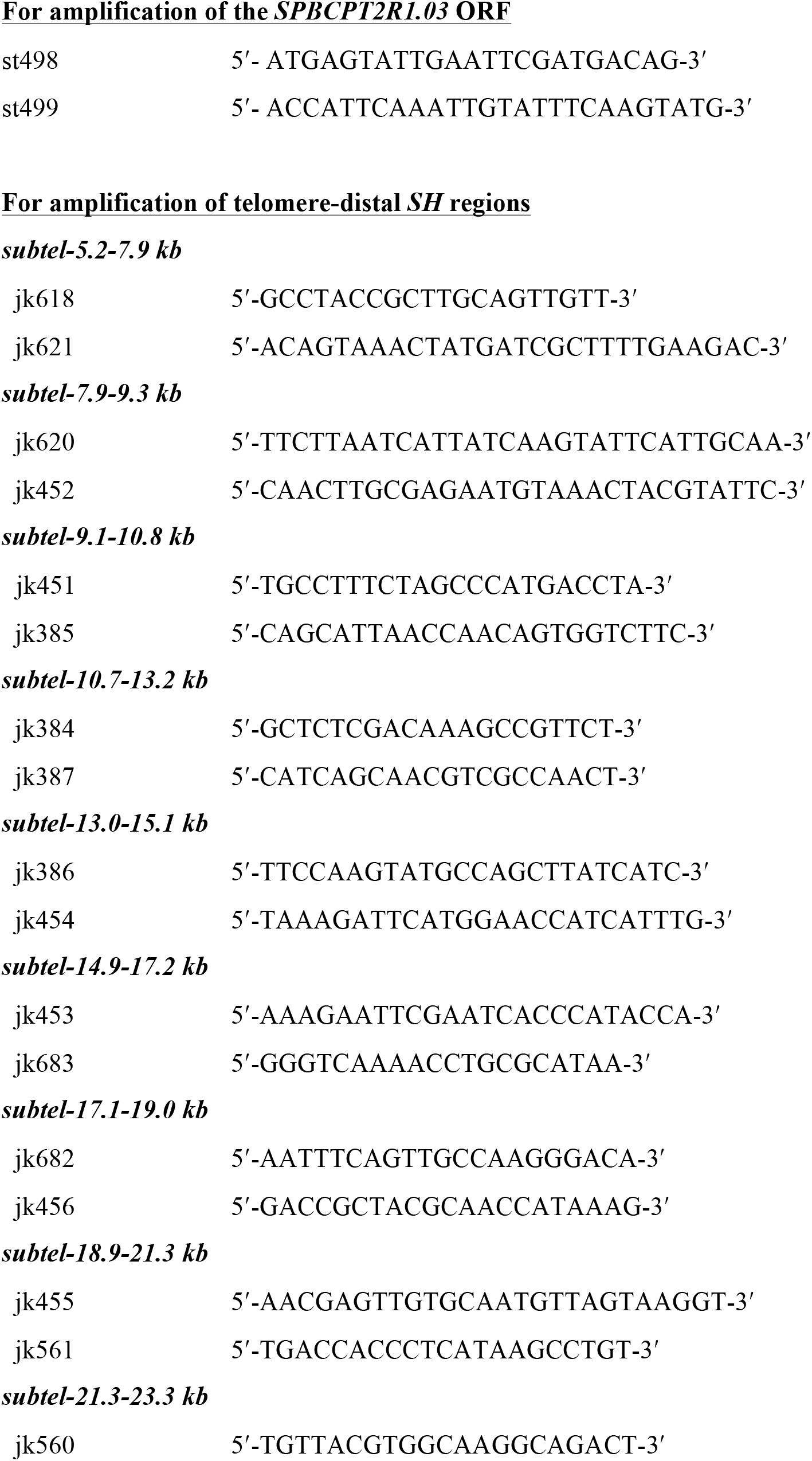
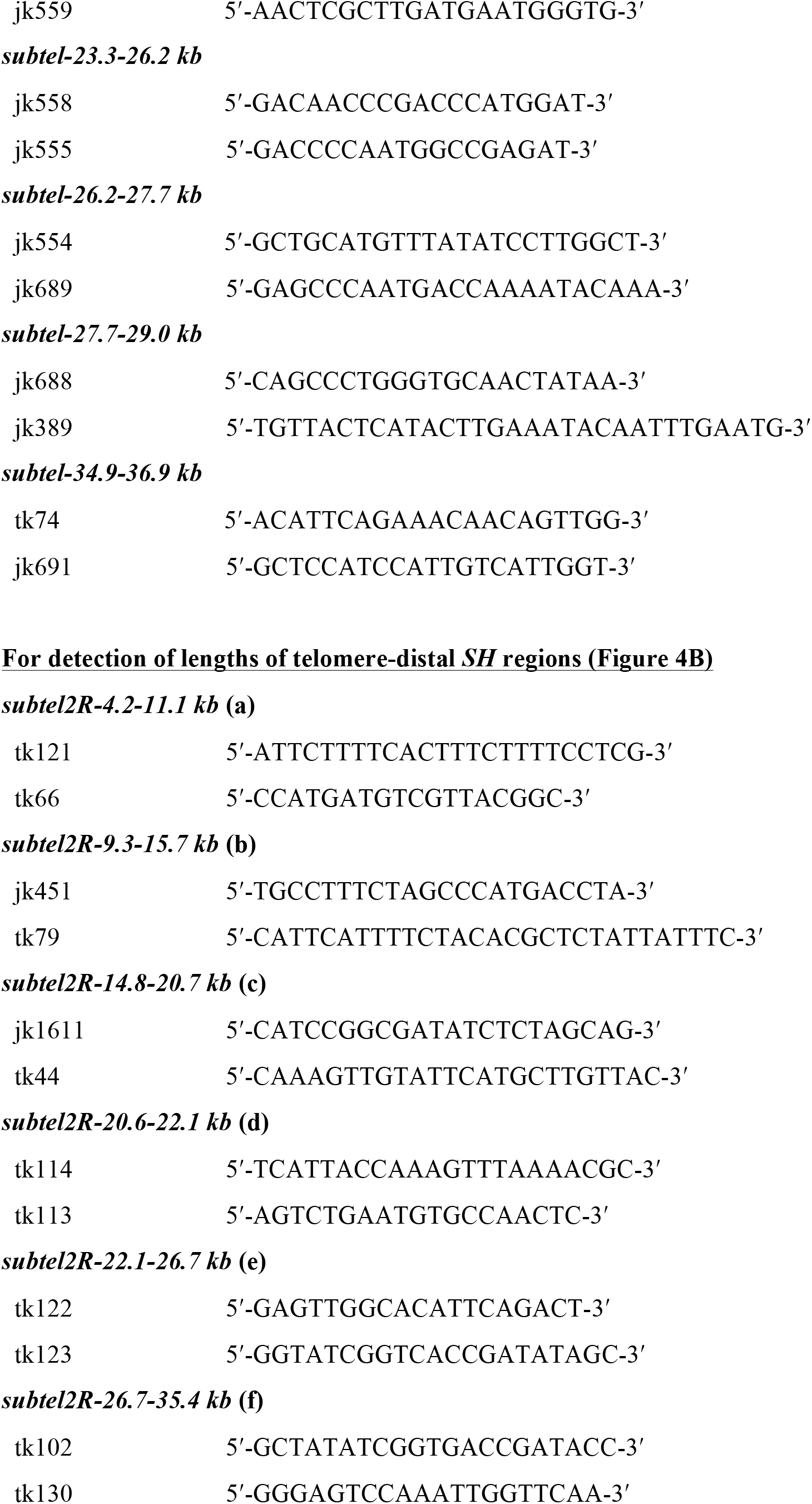
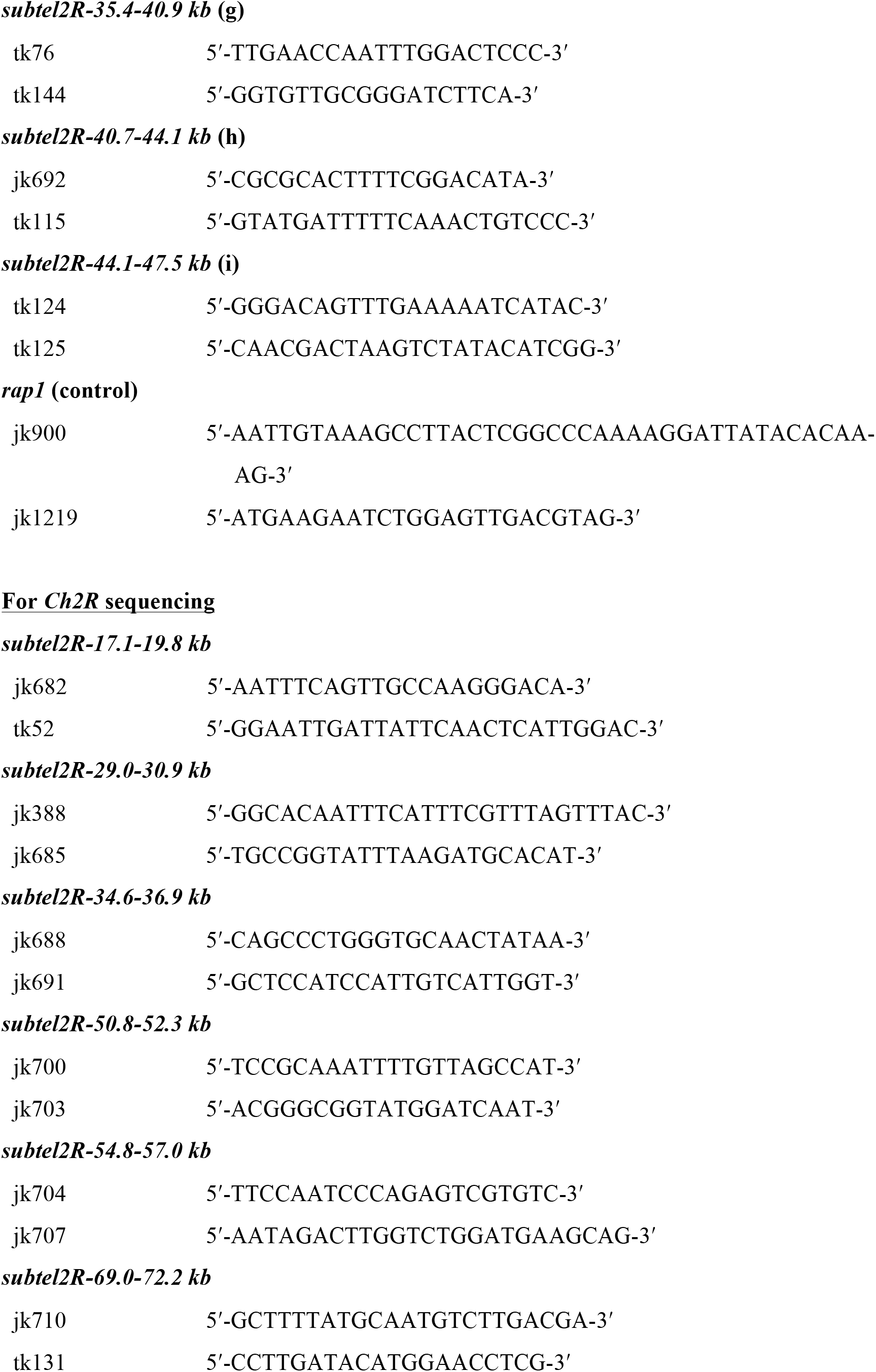
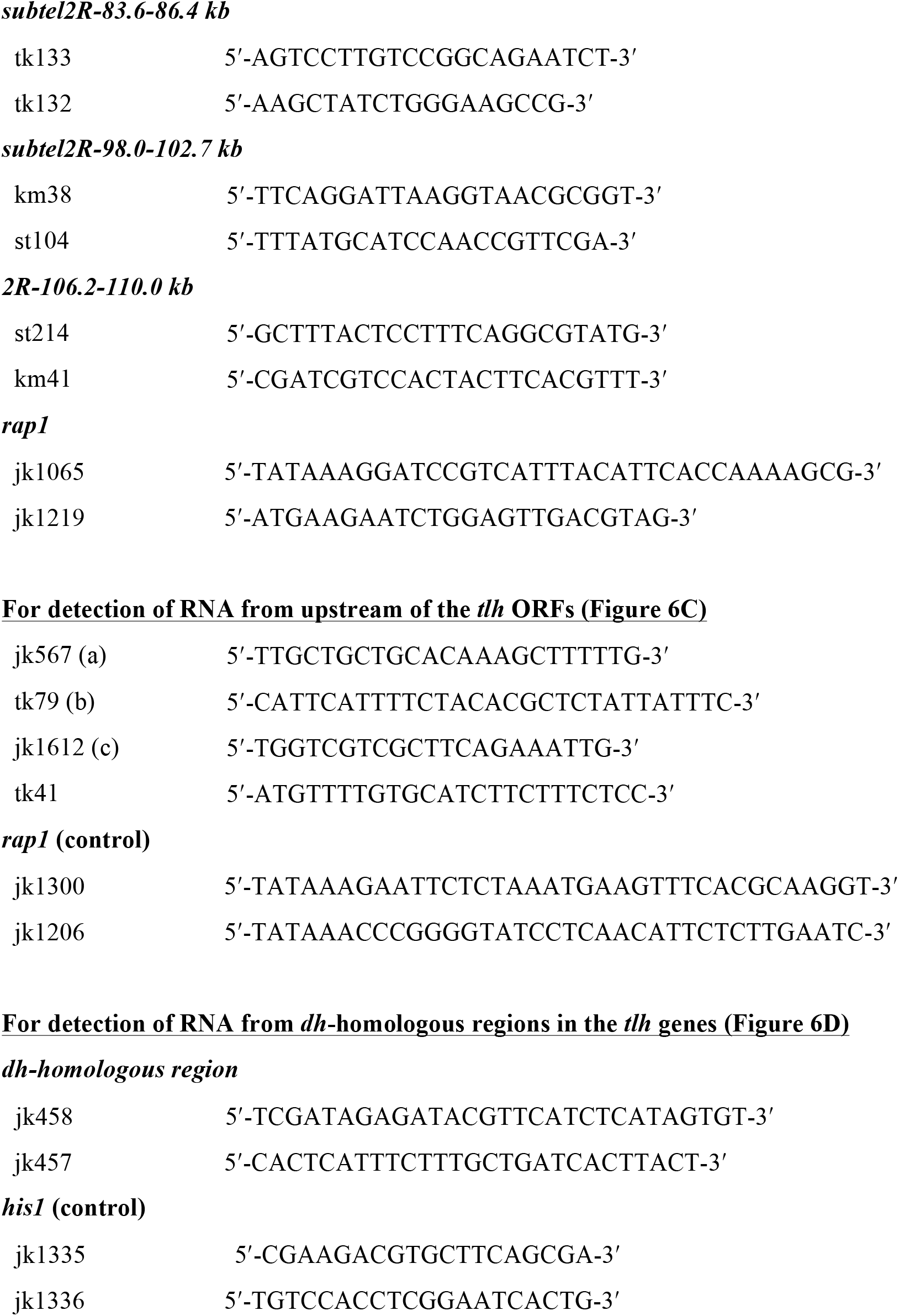
PCR primers used in this study.

**Table S3.**
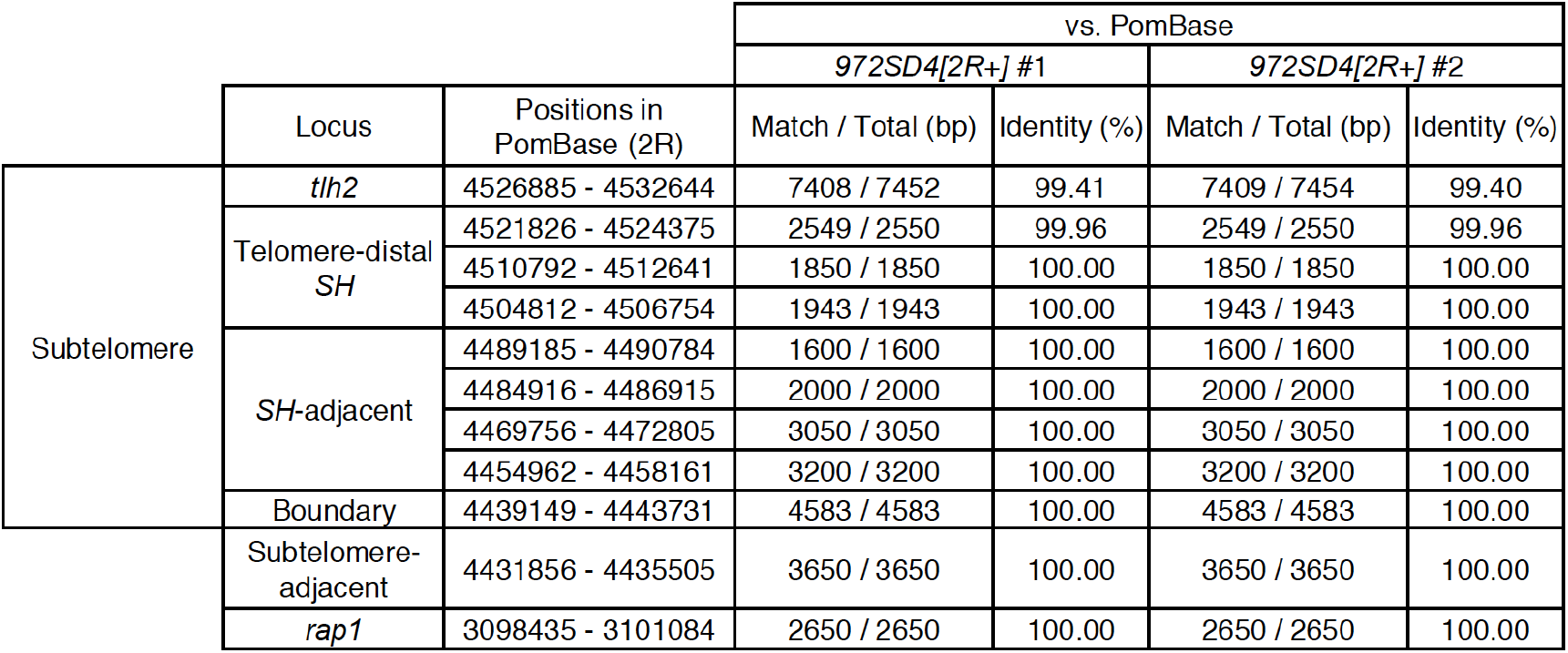
Sequence identities between sequences of *2R* in *972SD4[2R*+] (#1 and #2) and those in PomBase.

