## Supplemental Figures and Tables for "Complete sequences of *Schizosaccharomyces pombe* subtelomeres reveal multiple patterns of genome variation"

### for

and Junko Kanoh[1]*

[1] Institute for Protein Research, Osaka University, 3-2 Yamadaoka, Suita, Osaka, Japan.

[2] Present address: Institute of Molecular Biology, University of Oregon, 1370 Franklin Blvd, Eugene, OR, USA.

[3] These authors contributed equally to this work.

Contents: Figure S1-3

Tables S1–3

#### Supplementary Figures

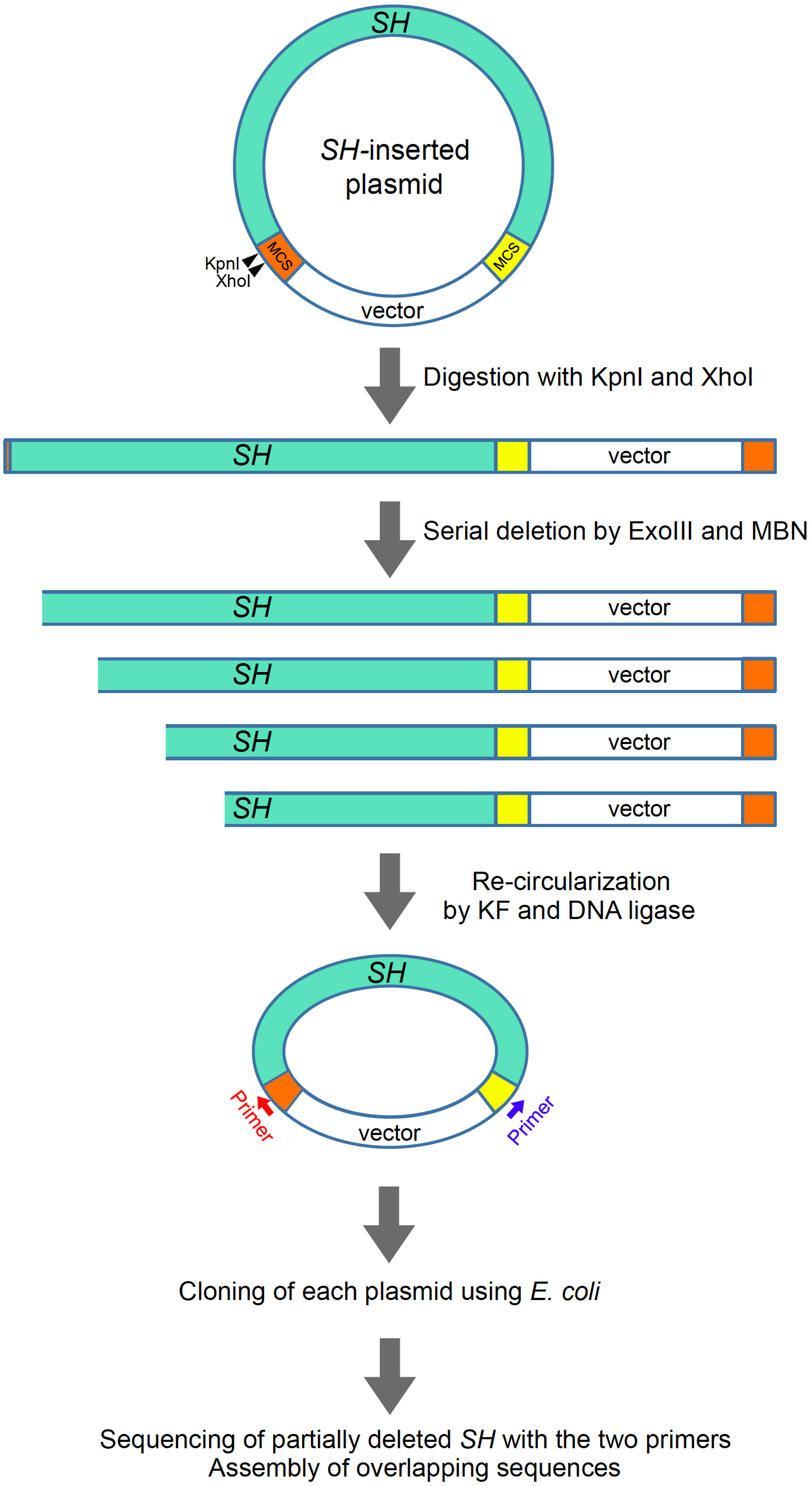

**Figure S1. Sequencing of telomere-proximal *SH* regions by the serial deletion method.**

A telomere-proximal *SH* region with various common segments (~5 kb) was amplified by PCR from each *972SD4* strain, and inserted into a vector. The resultant plasmid was digested with restriction enzymes, KpnI and XhoI, at the multi-cloning sites (MCSs) of the vector. The linearized plasmid was serially deleted from the 5′-protruding XhoI cutting site by treatment with exonuclease III (ExoIII) and mung bean nuclease (MBN) for fixed times. After deletion of the *SH* region, both ends of the plasmid were blunted by klenow fragment (KF) and ligated by DNA ligase. The resultant re-cirsularized plasmids were cloned using *E. coli*. The partially deleted *SH* region was sequenced using primers that aneal to MCSs of the vector. And the sequences were assembled using overlapping sequences.

**A**

```
A1   1  GCAGCCTCGCCTTACGGCTCGGCTGACGGGTGGGGCCCAATAGTGGGGGCATTGTATTTGTGAAAAAAAA   70
A2   1  GAATACTCGCCGTACGGCTCGGCTGACGGGTGGGGCCCAATAGTGGGGGCATTGTATTTGTGAAAAAAAA   70

    71  CTTAAGTAAATTTATTTTTTTATGAATACCGTATTTCATTTCTATTTCTTTATTCAACTTACCGCACTTC  140
    71  CTTAAGTAAATTTATTTTTTTATGAATACCGTATTTCATTTCTATTTCTTTATTCAACTTACCGCACTTC  140

   141  CTAAATTCCCTTTATTCTATATTGTTACTCCGTTACCCAATTAATTATGACTGAGTGAACTTCAAATTTT  210
   141  CTAAATTCCCTTTATTCTATATTGTTACTCCGTTACCCAATTAATTATGACTGAGTGAACTTCAAATTTT  210

   211  TCATCCATTTTAATTATCATAATACACTGC  240 (bp)
   211  TCATCCATTTTAATTATCATAATACACTGC  240 (bp)
```

A1 vs. A2
98.33%
(236 / 240 bp)

**B**

```
B    1  ACTACTGTATTACACTTAGTGAATTTCACCAT  32 (bp)
```

**C**

```
C1   1  ACTCAATTCCACACATTTCAATTTTTTCCCATCCTAAATACTATAGTACACTACCTCACCGTATTATACT   70
C2   1  ACTCAATTCCACACATTTCAATTTTTTCCCATCCTAAATACTATAGTACACTACCTCACCGTATTATACT   70
C3   1  ACTCAATTCCACACATTTCAATTTTTTCCCATCCTAAATACTATAGTACACTACCTCACCGTATTATACT   70
C'      ----------------------------------------------------------------------

    71  TAATGAACTCTAACACACTCAATTCATACAACTTCAAATTTTTTCCCATCCTAAATACTATAGTACACTA  140
    71  TAATGAACTCTAACACACTCAATTCATACAACTTCAAATTTTTTCCCATCCTAAATACTATAGTACACTA  140
    71  TAATGAACTCTAACACACTCAATTCATACAACTTCAAATTTTTTCCCATCCTAAATACTATAGTACACTA  140
        ----------------------------------------------------------------------

   141  CCTCACCGTATTATACTTAATGAACTCTAACACACTCAATTC–CACACATTTC-AATTTTTTACCCATCC  208
   141  CCTCACCGTATTATACTTAATGAACTCTAACACACTCAATTCATACAACTT-CAAATTTTTTACCTATCC  209
   141  CCTCACCGTATTATACTTAATGAACTCTAACACACTCAATTCATACAACTT-CAAATTTTTTACCTATCC  209
     1  ---------------------------------ACTCAATTC-CACACATTTC-AATTTTTTACCCATCC   35

   209  TAAATACTATAGTACACTACATCACCGTATTATACTTAATGAACTATAACACACTCAATTCATACAACTT  278
   210  TAAATACTATAGTACACTACATCACCGTATTATACTTAATGAACTCTAACACACTCAATTCATACAACTT  279
   210  TAAATACTATAGTACACTACATCACCGTATTATACTTAATGAACTCTAACACACTCAATTCATACAACTT  279
    36  TAAATACTATAGTACACTACATCACCGTATTATACTTAATGAACTATAACACACTCAATTCATACAACTT  105

   279  CAAATTTTTTACCCATCCTAATTATTATAGTACACTACATCACTGTATTATACTTAACGTATTCCAGCAC  348
   280  CAAATTTTTTAACCATCCTAATTATTATAGTACACTACCTCACCGTATTATACTTAACGTACTCCAGCAC  349
   280  CAAATTTTTTAACCATCCTAATTATTATAGTACACTACCTCACCGTATTATACTTAACGTACTCCAGCAC  349
   106  CAAATTTTTTACCCATCCTAATTATTATAGTACACTACATCACTGTATTATACTTAACGTATTCCAGCAC  175

   349  ACTCAATTCATACAACTTCAAATTTTTTATCTATCCTAAATACTATAGTACATTACATCACCGTATTATA  418
   350  ACTCAATTCC-ACACATTTCAATTTTTTACCCATCCTAAATACTATAGTACACTACATCACCGTATTATA  418
   350  ACTCAATTCATACAACTTCAAATTTTTTATCTATCCTAAATACTATAGTACATTACATCACCGTATTATA  419
   176  ACTCAATTCATACAACTTCAAATTTTTTATCTATCCTAAATACTATAGTACATTACATCACCGTATTATA  245

   419  CTTAATGAACTTTAACAC  436 (bp)
   419  CTTAATGAACTTTAACAC  436 (bp)
   420  CTTAATGAACTTTAACAC  437 (bp)
   246  CTTAATGAACTTTAACAC  263 (bp)
```

| C1 vs. C2 | C1 vs. C3 | C2 vs. C3 |
|---|---|---|
| 95.21% | 97.26% | 97.94% |
| (417 / 438 bp) | (426 / 438 bp) | (428 / 437 bp) |

## C*

```
c1  1  ACTCAATTCC-ACACATTTCAA-TTTTTT-CCCATCCTAAATACTATAGTACACTACCTCACCGTATTAT  67
c2  1  ACTCAATTCATACAACTT-CAAATTTTTT-CCCATCCTAAATACTATAGTACACTACCTCACCGTATTAT  68
c3  1  ACTCAATTCC-ACACATTTCAA-TTTTTTACCCATCCTAAATACTATAGTACACTACATCACCGTATTAT  68
c4  1  ACTCAATTCATACAACTT-CAAATTTTTTACCCATCCTAATTATTATAGTACACTACATCACTGTATTAT  69
c5  1  ACTCAATTCATACAACTT-CAAATTTTTTATCTATCCTAAATACTATAGTACATTACATCACCGTATTAT  69
c6  1  ACTCAATTCATACAACTT-CAAATTTTTTACCTATCCTAAATACTATAGTACACTACATCACCGTATTAT  69
c7  1  ACTCAATTCATACAACTT-CAAATTTTTTAACCATCCTAATTATTATAGTACACTACCTCACCGTATTAT  69
c8  1  ACTCAATTCC-ACACATTTCAA-TTTTTTACCCATCCTAAATACTATAGTACACTACATCACCGTATTAT  68

   68  ACTTAATGAACTCTAACAC  86 (bp)
   69  ACTTAATGAACTCTAACAC  87 (bp)
   69  ACTTAATGAACTATAACAC  87 (bp)
   70  ACTTAACGTATTCCAGCAC  88 (bp)
   70  ACTTAATGAACTTTAACAC  88 (bp)
   70  ACTTAATGAACTCTAACAC  88 (bp)
   70  ACTTAACGTACTCCAGCAC  88 (bp)
   69  ACTTAATGAACTTTAACAC  87 (bp)
```

| | c2 | | c3 | | c4 | | c5 | | c6 | | c7 | | c8 | |
|---|---|---|---|---|---|---|---|---|---|---|---|---|---|---|
| | Match /Total (bp) | Identity (%) | Match /Total (bp) | Identity (%) | Match /Total (bp) | Identity (%) | Match /Total (bp) | Identity (%) | Match /Total (bp) | Identity (%) | Match /Total (bp) | Identity (%) | Match /Total (bp) | Identity (%) |
| c1 | 82 / 88 | 93.18 | 84 / 87 | 96.55 | 73 / 89 | 82.02 | 77 / 89 | 86.52 | 80 / 89 | 89.89 | 75 / 89 | 84.27 | 84 / 87 | 96.55 |
| c2 | | | 80 / 89 | 89.89 | 78 / 88 | 88.64 | 82 / 88 | 93.18 | 85 / 88 | 96.59 | 80 / 88 | 90.91 | 80 / 89 | 89.89 |
| c3 | | | | | 74 / 89 | 83.15 | 79 / 89 | 88.76 | 81 / 89 | 91.01 | 74 / 89 | 83.15 | 86 / 87 | 98.85 |
| c4 | | | | | | | 77 / 88 | 87.50 | 79 / 88 | 89.77 | 84 / 88 | 95.45 | 74 / 89 | 83.15 |
| c5 | | | | | | | | | 85 / 88 | 96.59 | 78 / 88 | 88.64 | 80 / 89 | 89.89 |
| c6 | | | | | | | | | | | 79 / 88 | 89.77 | 82 / 89 | 92.13 |
| c7 | | | | | | | | | | | | | 74 / 89 | 83.15 |

## D

```
D1   1  ACTCAGTTCAAATCAACTTCAAATTTTTCATCCGTTTTATTCACCCATGCACACTACACCACCGTATTAC  70
D2   1  ACTCAGTTCAAATCAACTTCAAATTTTTCATCCGTTTTATTCACCCATACACACTACACCACCTTATTAC  70
D3   1  ACTCAGTTCAAATCAACTTCAAATTTTTCATCCGTTTTATTCACCCATACACACTACACCACCTTATTAC  70

    71  AATTAGTGAATTCCAACATAGTCTAATTCCATGAACTTCAACTTTTTGTCCATCTTCACTATTGCACTAC  140
    71  AATTAGTGAATTCCAACATAGTCTAATTCCATGAACTTCAACTTTTTGTCCATCTTCACTATTGCACTAC  140
    71  AATTAGTGAATTCCAACATAGTCTAATTCCATGAACTTCAACTTTTTGTCCATCTTCACTATTGCACTAC  140

   141  CCTACATCCTCGTATTCTACTTAATCTACTCCAACACCTTTTAATCCATTCATCTTCTTTTCTTTTACTC  210
   141  CCTACATCCTCGTATTCTACTTAATCTACTCCAACACCTTTTAATCCATTCATCTTCTTTTCTTTTACTC  210
   141  CCTACATCCTCGTATTCTACTTAATCTACTCCAACACCTTTTAATCCATTCATCTTCTTTTCTTTTACTC  210

   211  TTTTATTTTATTTATTTTATTTATTCTATTTATTCTATT  249 (bp)
   211  TTTTATTTTATTTATTCTATTTATTCTATTTATTCTATT  249 (bp)
   211  TTTTATTTTATTTATTTTATTTATTCTATTTATTCTATT  249 (bp)
```

| D1 vs. D2 | D1 vs. D3 | D2 vs. D3 |
|---|---|---|
| 98.89% | 99.20% | 99.60% |
| (246 / 249 bp) | (247 / 249 bp) | (248 / 249 bp) |

## E

```
E    1  ---------------------------------------------------------------TATTCTA  7
E'   1  ----------------------------------------CTATTTATTC----TATTCTATTTATTCTA  26
E"   1  TATTCTATTTATTCTATTTATTCTATTTATTCTATTTATTCTATTTATTCTATTTATTCTATTTATTCTA  70

     8  TTCCATTACTTTCTTCCTCTTCTTTCTTCCATAC------------------------------------  41
    27  TTCCATTACTTTCTTCCTCTTCTTTCTTCCATACTCTATCCTACCTACTCTCTACCTACTCTACCTACTC  96
    71  TTCCATTACTTTCTTCCTCTTCTTTCTTCCATACTCTATCCTACCTACTCTCTACCTACTCTACCTACTC  140

    42  --CCTACCTACTCTACCTATTCTC-------------------CTCCTTCATTTCCTTCT  92 (bp)
    97  TACCTACCTACTCTACCTATTCTCCTCCTTCATTTCCTTCTCTCCTTCATTTCCTTCT  166 (bp)
   141  TACCTACCTACTCTACCTATTCTC------------------------------------  176 (bp)
```

**F**

```
F1   1 CTCCTTCATTTCCTTCTCTCTCTCTAAAAACGTTTAACGAATACTCGCCTTACGGCTCGGCTGACGGGTG  70
F2   1 CTCCTTCATTTCCTTCTCTCTCTCTAAAAACGTTTAACGAATACTCGCCTTACGGCTCGGCTGACGGGTG  70
F3   1 CTCCTTCATTTCCTTCTCTCTCTCTAAAAACGTTTAACGAATACTCGCCTTACGGCTCGGCTGACGGGTG  70

    71 GGGG-CCAATAGTGGGGGCATTTTATATATGTGAAAAAAACTTAAGTAAATTTAAATTTTTATAAATAAC 139
    71 GGGC-CCAATAGTGGGGGCA--TTGTATATGTGAAAAAAACTTAAGTAAATTTAAATTTTTATAAATAAC 137
    71 GGGGTCCAGTAGTGGGGGCA--TTGTATATGTGAAAAAAACTTAAGTAAATTTAAATTTTTATAAATAAC 138

   140 GCAACTTAATTATCTTTCTTTATCT  164 (bp)
   138 GCAGCTTAATTATCTTTCTTTATCT  162 (bp)
   139 GCAGCTTAATTATCTTTCTTTATCT  163 (bp)
```

| F1 vs. F2 | F1 vs. F3 | F2 vs. F3 |
|---|---|---|
| 96.95% | 96.36% | 98.16% |
| (159 / 164 bp) | (159 / 165 bp) | (160 / 163 bp) |

**G**

```
G    1 ATATTACCGTACGTTCTAATTGCTCCTTATTCTACATTGTTACTTTGTAACCCAATTAATTATGACTGAG  70
    71 TGAACTTCAAATTTTTCATCCATTACCATTACCATATTACGCTGCACTACAGTATTACACTTAGTGAATT 140
   141 TCAACAT  147 (bp)
```

**H**

```
H1   1 TCATTATCATACTACGCTGCACAACTGTAATCTACTTAATCTACTCCAATATAGTCCTCTGCTTTTATCT  70
H2   1 TCATTATCATACTACGCTGCACAACTATAATCTACTTAATGTACTCCAATATAGTCCTCTGCTTTTATCT  70

    71 TCTTTCCTCTTACTTTACTTATTTCTTTTACTCTCTTACTTTATTTATTTACTCTCCTTACTCTGTTTCT 140
    71 TCTTTCCTCTTACTTTACTTATTTCTTTTACTCTCTTACTTTATTTATTTACTCTCCTTACTCTGTTTCT 140

   141 TCTATTCTATTCATCTTACTTCTTCCTCATTTCTTCTCTCTTCTCTTCTATATCTTCTACTCTTTCATAC 210
   141 TCTATTCTATTCATCTTACTTCTTCCTCATTTCTTCTCTCTTCTCTTCTATATCTTCTACTCTTTCATAC 210

   211 CTCTCTTACCTCCATCCATTATCCATTCTCTCTCTCCAAAATCGTTTAAC  260 (bp)
   211 CTCTCTTACCTCCATCCATTATCCATTCTCTCTCTCCAAAATCGTTTAAC  260 (bp)
```

| H1 vs. H2 |
|---|
| 99.23% |
| (258 / 260 bp) |

**I**

```
I    1 GCCCCTTTATTCTACTTACTGCACCCTAACGC  32 (bp)
```

**J**

```
J    1 ACTCAAGCCCACACATTTCAACTTTTTATTCACCATATCTCATTATATCTCCTAATCATACATAATGAAT  70
    71 CCTACCACGCTCAATTCAACCCAACTTCATTTTCTTCATTCATTTTCATTACCATAGTACGCTCCACAAC 140
   141 TGTAATCTACTTAATGTACTCCAACATACTCAACTTCCACACATTTCAAAATTTTTTTACCCATTTTAT 210
   211 TTACCCCTATTATACTTACCGTACTCCAACACACTCAATACATACCAACTTCAAATTTTTTATGCATCTG 280
   281 ACTTTCCAATATACACTATGTATCCATCCTTATTTACTTTACTTTCTAATACTCTCTTACTT  342 (bp)
```

**K**

```
K    1 TAATTACTTCTTTATTCATCTTATTCTATTTATTTTATTTATTCCTTACTCCCTACTCCCTACTCCCTAC  70
K'   1 TAATTACTTCTTTATTCATCTTATTCTATTTATTTTATTTATTCCT-------TACTCCCTACTCCCTAC  63

    71 TCCCTACTCTCTACTCTATTCTGTATTCTATATTCTATATTCTATATTCTATATTCTATATTCTATATTC 140
    64 TCCCTACTCTCTACTCTATTCTG--------------TATTCTATATTCTATATTCTATATTCTATATTC 119

   141 TATATTCTATATTCTATATTTTCTTTTACCTCCTTTATCTTTCGTTAACATGTTTAACGAATACTCGCTT 210
   120 TATATTCTATATTCTATATTTTCTTTTACCTCCTTTATCTTTCGTTAACATGTTTAACGAATACTCGCTT 189

   211 TACAGCTCGTTACCCACCTCACACATCACTGACTCAACTTCAAATTTCTCAACCGTACTCATTACCATAG 280
   190 TACAGCTCGTTACCCACCTCACACATCACTGACTCAACTTCAAATTTCTCAACCGTACTCATTACCATAG 259

   281 TACGCTGCGCAACTGTAATCTACTTAATGAACCCTAA  317 (bp)
   260 TACGCTGCGCAACTGTAATCTACTTAATGAACCCTAA  296 (bp)
```

**L**

```
L1   1  CACACTCAATTCAAATCAACTTCAAATTTTTCATTCGTTTTATTCACCCATG  52 (bp)
L2   1  CACACTCAATTCAAATCAACTTCAAATTTTTCATCCGTTTTATTCACTCATG  52 (bp)
```

L1 vs. L2
96.15%
(50 / 52 bp)

**M**

```
M    1  TACACTATACTACCTTATTACCCTTAGTGAACCATAACACACTCACTTCAACCCAACTTCAAATTTTTCA  70
    71  TCCGTTTTATTCCACTCATGTACACTACACCACCTTATTCCTACACATTTTTATTCATTTTACTTTCTTA  140
   141  TTCTCCTACTCTTCTTATTCCATTATTTTCTTCCTCTTCTTTCTTCCATACTCTACTCTATCTGTTCTAC  210
   211  CTATTCTCCTCCTTCCTTCTCTCCTTTATTCTGCCCCTTCATTTTCTCCTTCATTTCCTTATACTTCTCT  280
   281  AACTACG  287 (bp)
```

**N**

```
N    1  TGATTACTTCTTCATTCTCCTACTTTATTTATTTTATTCATTTTATTTTATTTTATTTTATTTATTCGTT  70
    71  TTTAAATATTTAATTTACTCTCCTTACTTTATTTTTTTGATTTCTACTCTCTTCTATATATCCTACTCTT  140
   141  ATATTCTCTCCTTCATTTCCTTATTCCATATTGTTACTTCGTTATCCAGCTCACCATGACTGAGTCAACT  210
   211  TCAAATTTTTCATCCCCTTCAATTACCAAAGTCCACTACAATTCTCTATTTATATTCCTGCATCCCAACA  280
   281  CATTATTTCAACCACTTGTCTATCATAATTACTATAGTTCATTACTCATCCCTATTCTACTTAATGCACT  351
   351  CCAATATAGTCCCCTGCTTTTATCTTCTTTCCTCTTACTTTAATTGCTTCTTTATTCTCCTACCTTATTT  421
   421  ATTTTATTTATTTTATTTATTTTATTTTATTTTATTTATTTTATTTTACTCTCCTTACTCTATTTCTTCT  491
   491  ATATATTCTACTTATATATCCTGCTCCTTCATCCTACTCTCTTCTCTCTATATTTTACTCTTAAATTCTA  561
   561  TCTGTTACATTCTTGTCTTTCGTTAACAACA  591 (bp)
```

**O**

```
O1   1  TTTAACGATTACTCGCCTTACGGCTCGTTACTTGCTTCACACCTGACTAAGTGAACTTTGAATTTTTTTT  70
O2   1  TTTAACGATTACTCGCCTTACGTCTCGTTGCTCGCTTCACACCTGACTAAGTGAACTTTGAATTTTTTTT  70

    71  CCTTTATTAAAATGACCATCCTCAAAATTTCCCATCCTC  109 (bp)
    71  CCTTTATTAAAATGACCATCCTCAAAATTTCCCATCCTC  109 (bp)
```

O1 vs. O2
97.25%
(106 / 109 bp)

**P**

```
P1   1  CCCGACGACATATCACTCAACAAACGTTTTCAATAATTTTTTTTTCTTTTATCAATCATTTTATAATTTA  70
P2   1  CCCGACGACATATCACTCAACAAACTTTTTCAATAATTTTTTTTTCTTTTATTAATCATTTTATAATTTA  70
P3   1  CCCGACGACATATCACTCAACAAACTTTTTCAATAATTTTTTTTTCTTTTATTAATCATTTTATAATTTA  70
P4   1  CCCGACGACATATCACTCAACAAACGTTTTCAATAATTTTTTTTTCTTTTATCAATCATTTTATGATTTA  70

    71  TTTTCAATTTTCAACTGCCTGCTGTTTGTCGTTCATTGGCAAATAAATATTTTTCATTATTTTCTATCAT  140
    71  TTTTCAATTTTCAACTGCCTGCTGTTTGTCATTCATTGGCAAATAAATATTTTTCATTATTTTCTATCAT  140
    71  TTTTCAATTTTCAACTGCCTGCTGTTTGTCGTTCATTGGCAAATAAATATTTTTCATTATTTTCTATCAT  140
    71  TTTTCAATTTTCAACTGCCTGCTGTTTGTCGTTCATTGGCAAATAATTATTTTTCATTATTTTCTATCAT  140

   141  TATTTAATATTGTCTTTATTTGTGTTCCAATATTTGTATTTATATTTTTTATTTCATTTTATTTTG  206 (bp)
   141  TATTTAATATTGTCTTTATTTGTGTTCCAATATTTGTATTTATATTTTTTATTTCATTTTATTTTG  206 (bp)
   141  TATTTAATATTGTCTTTATTTGTGTTCCAATATTTGTATTTATATTTTTTATTTCATTTTATTTTG  206 (bp)
   141  TATTTAATATTGTCTTTATTTGTGTTCCAATATTTGTATTTATATTTTTTATTTCATTTTATTTTG  206 (bp)
```

| | P2 | | P3 | | P4 | |
|---|---|---|---|---|---|---|
| | Match /Total (bp) | Identity (%) | Match /Total (bp) | Identity (%) | Match /Total (bp) | Identity (%) |
| P1 | 203 / 206 | 98.54 | 204 / 206 | 99.03 | 204 / 206 | 99.03 |
| P2 | | | 205 / 206 | 99.51 | 201 / 206 | 97.57 |
| P3 | | | | | 202 / 206 | 98.06 |

# Q

```
Q1  1  TTAATTTTATTTATTTCTGTATATTTATTCGTTTTTAAATATTTTATTTTATTTTATTTCATTTCATTTC  70
Q2  1  TTAATTTTATTTATTTCTATATATTTATTCGTTTTTAAATATTTTATTTTATTTTATTTCATTTCATTTC  70
Q3  1  TTAATTTTATTTATTTCTATATATTTATTCGTTTTTAAATATTTTATTTTATTTT-----ATTTCATTTC  65
Q4  1  TTAATTTTATTTATTTCTGTATATTTATTCGTTTTTAAATATTTTATTTTATTTTTATTTCATTTC  70
Q5  1  TTAATTTTATTTATTTCTATATATTTATTCGTTTTTAAATATTTTATTTTATTTTATTTCATTTCATTTC  70

   71  ATTTTATTTTTTTATTTATTTTACACAATCATATTATTTGTTTAACATGACTAACATAAGCTCAATTTTC  140
   71  ATTTTATTTTTTTATTTATTTTACACAATCATATTATTTGTTTAACATGACTAACATAAGCTCAATTTTC  140
   66  ATTTTATTTTTTTATTTATTTTACGCAATCATATTATTTATTTAACATATCTAACATAAGCTCAATTTTC  135
   71  ATTTTATTTTTTTATTTATTTTACACAATCATATTATTTGTTTAACATGACTAACATAAGCTCAATTTTC  140
   71  ATTTTATTTTTTTATTTATTTTACGCAATCATATTATTTATTTAACATATCTAACATAAGCTCAATTTTC  140

  141  AAACACTTCATTTATCAAAGTACACCGCATGTTCCTATTATACTTACTGCACTCCAACATA  201 (bp)
  141  AAACACTTCATTTATCAAAGTACACCGCATGTTCCTATTATACTTACTGCACTCCAACATA  201 (bp)
  136  AAACACTTCATTTATCAAAGTACACCGCATGTTCCTATTATACTTACTGCACTCCAACATA  196 (bp)
  141  AAACACTTCATTTATCAAAGTACACCGCATGTTCCTATTATACTTACTGCACTCCAACATA  201 (bp)
  141  AAACACTTCATTTATCAAAGTACACCGCATGTTCCTATTATACTTACTGCACTCCAACATA  201 (bp)
```

| | Q2 | | Q3 | | Q4 | | Q5 | |
|---|---|---|---|---|---|---|---|---|
| | Match /Total (bp) | Identity (%) | Match /Total (bp) | Identity (%) | Match /Total (bp) | Identity (%) | Match /Total (bp) | Identity (%) |
| Q1 | 200 / 201 | 99.50 | 191 / 201 | 95.02 | 200 / 201 | 99.50 | 196 / 201 | 97.51 |
| Q2 | | | 192 / 201 | 95.52 | 199 / 201 | 99.00 | 197 / 201 | 98.01 |
| Q3 | | | | | 191 / 201 | 95.02 | 196 / 201 | 97.51 |
| Q4 | | | | | | | 195 / 201 | 97.01 |

# R

```
R1  1  GTCCCCTTCATTGTATCTTCTTTTTTCTTACTTTACTTACTTCTTTGTTCTCCCATTTTTTATTATTATT  70
R2  1  GTCCCCTTCATTGTATCTTCTTTTTTCTTACTTTACTTACTTCTTTGTTCTCCCATTTTTTATTATTATT  70

   71  TCTTATTCCATTACTTTCTTCCTCTTCTTTCTTCCATATTCTAC  114 (bp)
   71  TCTTATTCCATTACTTTCTTCCTCTTCTTTCTTCCATACTCTAC  114 (bp)
```

R1 vs. R2
99.12%
(113 / 114 bp)

# S

```
S   1  TCTACCTATTCTGCTTTACTTATTCTACCTATTCTACCTATTCTGCTTTACTTATTCTACCTATCCAATA  70
S'  1  TCTACCTATTCTGCTTTACTTATTCTACCTAT--------------------------------CCAATA  38

   71  TTTATACTTCCTATGTTTTACCTCTTTTATATTTCGTTAACATGTTTAACGATTACTCGCTTTACTGCTC  140
   39  TTTATACTTCCTATGTTTTACCTCTTTTATATTTCGTTAACATGTTTAACGATTACTCGCTTTACTGCTC  119

  141  GTTACCCACCTGAATCAAAACTGAGTCAACTTCATATTTTT  181 (bp)
  120  GTTACCCACCTGAATCAAAACTGAGTCAACTTCATATTTTT  149 (bp)
```

# T

```
T   1  CATCCATCTTGATTATCATAATACACTGCGTATCTTTATTCAACTTACTGCATTACAACACAGCCTTATT  70
   71  CTATAAACTTCAAGTTTTCATTCATCTATCCTCATTCGCCATAT  114 (bp)
```

# U

```
U1  1  TGCACTACATCCCCTTATTATACTTACTGCATCCCAACACACTCACTTCCACTCATTTCAGTGTTTTGTC  70
U2  1  CACACTACATCCCCTTATTATACTTACTGCATCCCAACACACTCACTTCCACTCATTTCAGTGTTTTGTC  70
U3  1  TGCACTACATCCCCTTATTATACTTACTGCATCCCAACACACTCACTTCCACTCATTTCAGTGTTTTATC  70

   71  CATCTTCACTACCACACTACACTACTCATTCTT  103 (bp)
   71  CATCTTCACTACCACACTACACTACTCATTCTT  103 (bp)
   71  CATCTTCACTACCACACTACACTACTCATTCTT  103 (bp)
```

| V1 vs. V2 | V1 vs. V3 | V2 vs. V3 |
|---|---|---|
| 98.06% | 99.03% | 97.09% |
| (101 / 103 bp) | (102 / 103 bp) | (100 / 103 bp) |

**V**

V 1 TTAATTTTATTTCA 14 (bp)

**W**

W 1 TCTACTTATTCTACCTATACCTATTCTAC 29 (bp)

**X**

X 1 TAACCATCCTAATTATTATAGTACACTACTTGTCCCTATCACCTTTAACGTACTCCAG 58 (bp)

**Figure S2. Sequences of common segments in telomere-proximal *SH* regions.**

Sequences of subtypes and variants of each segment are aligned. Nucleotide changes are shown in red. Sequence identities between variants of each segment are indicated. Positions of common sequence motifs in segments C, E, K, and S are indicated according to Figure 3B. Panel C* shows sequences of the common sequence motifs of segment C, c1–8, and identities between them.

# A

```
SH1L(L) 1   ATTGATTGCTTTAATTTCAAACATGTCCCCTTTAATAGTTGGGACTTTAATAATTATCCTATTGTCAGGA  70
SH1L(R) 1   ATTGATTGCTTTAATTTCAAACATGTCCCCTTTAATAGTTGGGACTTTAATAATTATCCTATTGTCAGGA  70
SH1R    1   ATTGATTGCTTTAATTTCAAACATGTCCCCTTTAATAGTTGGGACTTTAATAATTATCCTATTGTCAGGA  70
SH2L    1   ATTGATTGCTTTAATTTCAAACATGTCCCCTTTAATAGTTGGGACTTTAATAATTATCCTATTGTCAGGA  70
SH2R    1   ATTGATTGCTTTAATTTCAAACATGTCCCCTTTAATAGTTGGGACTTTAATAATTATCCTATTGTCAGGA  70

        71  CTCGCAACTGCTTTTTATGTTACGTGGCAAGGCAGACTCATTTGTGCTGGTGTAGGGCTCATACTTGAAC  140
        71  CTCGCAACTGCTTTTTATGTTACGTGGCAAGGCAGACTCATTTGTGCTGGTGTAGGGCTCATACTTGAAC  140
        71  CTCGCAACTGCTTTTTATGTTACGTGGCAAGGCAGACTCATTTGTGCTGGTGTAGGGCTCATACTTGAAC  140
        71  CTCGCAACTGCTTTTTATGTTACGTGGCAAGGCAGACTCATTTGTGCTGGTGTAGGGCTCATACTTGAAC  140
        71  CTCGCAACTGCTTTTTATGTTACGTGGCAAGGCAGACTCATTTGTGCTGGTGTAGGGCTCATACTTGAAC  140

       141  AGGCTTATGAGGGTGGTCAGATGTTTAACACATTGATGGCACATTGCTTTGAAACGTACAATGGTGTTGA  210
       141  AGGCTTATGAGGGTGGTCAGATGTTTAACACATTGATGGCACATTGCTTTGAAACGTACAATGGTGTTGA  210
       141  AGGCTTATGAGGGTGGTCAGATGTTTAACACATTGATGGCACATTGCTTTGAAACGTACAATGGTGTTGA  210
       141  AGGCTTATGAGGGTGGTCAGATGTTTAACACATTGATGGCACATTGCTTTGAAACGTACAATGGTGTTGA  210
       141  AGGCTTATGAGGGTGGTCAGATGTTTAACACATTGATGGCACATTGCTTTGAAACGTACAATGGTGTTGA  210

       211  GAAAAGTGGAACGCAGTGTGTGGCCGATTGGCTTAAAGTAGGGCTTTTGGCTGTCACATTTGGGGCTGGA  280
       211  GAAAAGTGGAACGCAGTGTGTGGCCGATTGGCTTAAAGTAGGGCTTTTGGCTGTCACATTTGGGGCTGGA  280
       211  GAAAAGTGGAACGCAGTGTGTGGCCGATTGGCTTAAAGTAGGGCTTTTGGCTGTCACATTTGGGGCTGGA  280
       211  GAAAAGTGGAACGCAGTGTGTGGCCGATTGGCTTAAAGTAGGGCTTTTGGCTGTCACATTTGGGGCTGGA  280
       211  GAAAAGTGGAACGCAGTGTGTGGCCGATTGGCTTAAAGTAGGGCTTTTGGCTGTCACATTTGGGGCTGGA  280

       281  GGACCTAGATTGGTTAACACATTAGGTGGT  301 (bp)
       281  GGACCTAGATTGGTTAACACATTAGGTGGT  301 (bp)
       281  GGACCTAGATTGGTTAACACATTAGGTGGT  301 (bp)
       281  GGACCTAGATTGGTTAACACATTAGGTGGT  301 (bp)
       281  GGACCTAGATTGGTTAACACATTAGGTGGT  301 (bp)
```

# B

```
SH1L(L)              1 G-------------------AAAGCCAAAATTAAAATCTTGTAGACCAGCCAATGGCCGAAATATAAGAT 51
SH1L(R)              1 GTGTCACAATTTTAAAGCCTAAAGACAATATTACAAGTTTGTAAAGAGCCCAATGACCAAAATACAAAAG 70
SH1R                 1 GTGTCACAATTTTAAAGCCTAAAGACAATATTACAAGTTTGTAAAGAGCCCAATGACCAAAATACAAAAG 70
SH2L(L), SH2R(L)     1 GTGTCACAATTTTAAAGCCTAAAGACAATATTACAAGTTTGTAAAGAGCCCAATGACCAAAATACAAAAG 70
SH2L(R), SH2R(R)     1 GTGTCACAATTTTAAAGCCTAAAGACAATATTACAAGTTTGTAAAGAGCCCAATGACCAAAATACAAAAG 70

                    52 TCCAACCATAGCTGCTATAACTCCTGCTATTGTTGCAGTGATAATCACACTGAACGCTGCTATAACTCCT 121
                    71 GCCAGTTATAGTTG---------------------------------CACCCAGGGCTGCTATAATCATT 107
                    71 GCCAGTTATAGTTG---------------------------------CACCCAGGGCTGCTATAATCATT 107
                    71 GCCAGTTATAGTTG---------------------------------CACCCAGGGCTGCTATAATCATT 107
                    71 GCCAGTTATAGTTG---------------------------------CACCCAGGGCTGCTATAATCATT 107

                   122 GCTATTGTTGCAGTGATAATCACACTGAACGCTGCTATAACCACACCG--------ACCCA----TGCTA 179
                   107 CCCAAGGCTGC---TATAATCATTCCCAAGACTGCTATAATCATACCCAAGACTGCACCCAAGACTGCTA 174
                   108 CCCAAGGCTGC---TATAATCATTCCCAAGACTGCTATAATCATACCCAAGACTGCACCCAAGACTGCTA 174
                   108 CCCAAGGCTGC---TATAATCATTCCCAAGACTGCTATAATCATACCCAAGACTGCACCCAAGACTGCTA 174
                   108 CCCAAG------------------------ACTGCTATAATCATACCCAAGACTGC------------TA 141

                   180 TAACCATACTGATTAACGGTAGAGCTTTTATAATAAGCTTGGAGCAGTCTACAATAAACTATTAGTAAAG 249
                   175 TAATCACACCGGTCACCCGTAAAGCAAGTACCGCAAACTCTGCAAAGACTATTTTAAATTATTAGCAAAA 244
                   175 TAATCACACCGGTCACCCGTAAAGCAAGTACCGCAAACTCTGCAAAGACTATTTTAAATTATTAGCAAAA 244
                   175 TAATCACACCGGTCACCCGTAAAGCAAGTACCGCAAACTCTGCAAAGACTATTTTAAATTATTAGCAAAA 244
                   142 TAATCACACCGGTCACCCGTAAAGCAAGTACCGCAAACTCTGCAAAGACTATTTTAAATTATTAGCAAAA 211

                   250 AGA---AAAAACAGCTTCAGTGTGAAGGAAACGTACAGTTAAAAGCAGACAATAAGAGTATGAATAAGCC 316
                   245 AAAAAGAAAAAAAGCATCACTATAAAGGAAACATACAAATTAAACTAAGTAACATTAAAACAAGCAAACA 314
                   245 AAAAAGAAAAAAAGCATCACTATAAAGGAAACATACAAATTAAACTAAGTAACATTAAAACAAGCAAACA 314
                   245 AAAAAGAAAAAAAGCATCACTATAAAGGAAACATACAAATTAAACTAAGTAACATTAAAACAAGCAAACA 314
                   212 AAAAAGAAAAAAAGCATCACTATAAAGGAAACATACAAATTAAACTAAGTAACATTAAAACAAGCAAACA 281

                   317 AACATAAATAACAATAGGCCCCCAAATTTCACTGTGGGACTGATTTTCACCTATCCATTCTAAATATGAC 386
                   315 AACGTAAATAGCAATAGGCACCCAAATTTCACTGTAGAACCGATTTTCATTTATCCAAATTGAAAGAAAC 384
                   315 AACGTAAATAGCAATAGGCACCCAAATTTCACTGTAGAACCGATTTTCATTTATCCAAATTGAAAGAAAC 384
                   315 AACGTAAATAGCAATAGGCACCCAAATTTCACTGTAGAACCGATTTTCATTTATCCAAATTGAAAGAAAC 384
                   282 AACGTAAATAGCAATAGGCACCCAAATTTCACTGTAGAACCGATTTTCATTTATCCAAATTGAAAGAAAC 351

                   387 TCTTTTATTTTAAATTTCTTGTCTATTGTTAAAATAGTGATAATAGACCAAATAATAAGCAAGACCCATC 456
                   385 TTTTGGACTTCGAACTTTTTGTCTACTGCGAATACAATGATAATAGACCAAATAATAAGCAAGGGCCAAC 454
                   385 TTTTGGACTTCGAACTTTTTGTCTACTGCGAATACAATGATAATAGACCAAATAATAAGCAAGGGCCAAC 454
                   385 TTTTGGACTTCGAACTTTTTGTCTACTGCGAATACAATGATAATAGACCAAATAATAAGCAAGGGCCAAC 454
                   352 TTTTGGACTTCGAACTTTTTGTCTACTGCGAATACAATGATAATAGACCAAATAATAAGCAAGGGCCAAC 421

                   457 GAGATTTGTCCTGAGTAGAAATTCTTCCATTAGGATAATAGCTATTGTGGAATTTAGAAAATTCAGTAAA 526
                   455 GAAAATTACTGCGGGTAGAAATCCTTCCATCAGGATAATAGCTATTGTGGAATTTAGAAAATTCAGTAAA 524
                   455 GAAAATTACTGCGGGTAGAAATCCTTCCATCAGGATAATAGCTATTGTGGAATTTAGAAAATTCAGTAAA 524
                   455 GAAAATTACTGCGGGTAGAAATTCTTCCATCAGGATAATAGCTATTGTGGAATTTAGAAAATTCAGTAAA 524
                   422 GAAAATTACTGCGGGTAGAAATCCTTCCATCAGGATAATAGCTATTGTGGAATTTAGAAAATTCAGTAAA 491

                   527 AA 528 (bp)
                   525 AA 526 (bp)
                   525 AA 526 (bp)
                   525 AA 526 (bp)
                   492 AA 493 (bp)
```

# C

| | *SH1L*(R), *SH1R* | | *SH2L*(L), *SH2R*(L) | | *SH2L*(R), *SH2R*(R) | |
|---|---|---|---|---|---|---|
| | Match /Total (bp) | Identity (%) | Match /Total (bp) | Identity (%) | Match /Total (bp) | Identity (%) |
| *SH1L*(L) | 352 / 562 | 62.63 | 353 / 562 | 62.81 | 330 / 558 | 59.14 |
| *SH1L*(R), *SH1R* | | | 525 / 526 | 99.81 | 493 / 526 | 93.73 |
| *SH2L*(L), *SH2R*(L) | | | | | 492 / 526 | 93.54 |

**Figure S3. Homologous sequences at the ends of insertions in telomere-distal *SH* regions.**

(A) Homologous sequences at the ends of the 3.7 kb-insertion (pink boxes in Figure 4A). *SH1L* (L), a homologous sequence on the left side of the 3.7 kb-insertion in *SH1L* in Figure 4A; *SH1L* (R), that on the right side of the 3.7 kb-insertion in *SH1L*; *SH1R*, that in *SH1R*; *SH2L*, that in *SH2L*; *SH2R*, that in *SH2R*. Note that these sequences are 100% identical.

(B) Homologous sequences at the ends of the 7.1 kb-insertion (red, orange, and brown boxes in Figure 4A). Note that the region containing multiple repeat units (1–5) shows high sequence variation. Sequence variations among *SH1L* (R), *SH1R, SH2L* (L), *SH2R* (L), *SH2L* (R), and *SH2R* (R) are highlighted in orange.

(C) Sequence identities between homologous sequences in (B). Note that sequences of the pairs, *SH1L* (R) and *SH1R, SH2L* (L) and *SH2R* (L), and *SH2L* (R) and *SH2R* (R) are 100% identical.

#### Supplementary Tables

**Table S1. Fission yeast strains used in this study.**

| | |
|---|---|
| 972/JK107 | *h*$^-$ |
| JP1225 | *h*$^-$ *ade6-M216 leu1-32 ura4-D18 his7-366* (a derivative of CHP429) |
| ST3524 (*SD5*) | *h*$^+$ *ade6-M210 leu1-32 ura4-D18 his7-366 SH1L::ura4*$^+$ *SH1R::his7*$^+$ *SH2L::his7*$^+$ *SH2R::his7*$^+$ *SH3L::ura4*$^+$ |
| ST3479 (*SD5*) | *h*$^-$ *ade6-M216 leu1-32 ura4-D18 his7-366 SH1L::ura4*$^+$ *SH1R::his7*$^+$ *SH2L::his7*$^+$ *SH2R::his7*$^+$ *SH3L::ura4*$^+$ |
| ST5057 (*972SD4[1L+]* #1) | *h*$^+$ *ade6-M216 leu1-32 ura4-D18 his7-366 SH1R::his7*$^+$ *SH2L::his7*$^+$ *SH2R::his7*$^+$ *SH3L::ura4*$^+$ |
| ST5058 (*972SD4[1L+]* #2) | *h*$^-$ *ade6-M216 leu1-32 ura4-D18 his7-366 SH1R::his7*$^+$ *SH2L::his7*$^+$ *SH2R::his7*$^+$ *SH3L::ura4*$^+$ |
| ST5345 (*972SD4[1L+]* #3) | *h*$^-$ *ade6-M210 leu1-32 ura4-D18 SH1R::his7*$^+$ *SH2L::his7*$^+$ *SH2R::his7*$^+$ *SH3L::ura4*$^+$ |
| ST5059 (*972SD4[1R+]* #1) | *h*$^-$ *ura4-D18 SH1L::ura4*$^+$ *SH2L::his7*$^+$ *SH2R::his7*$^+$ *SH3L::ura4*$^+$ |
| ST5346 (*972SD4[1R+]* #2) | *h*$^+$ *ade6-M216 leu1-32 ura4-D18 his7-366 SH1L::ura4*$^+$ *SH2L::his7*$^+$ *SH2R::his7*$^+$ *SH3L::ura4*$^+$ |
| ST5347 (*972SD4[1R+]* #3) | *h*$^+$ *ade6-M210 leu1-32 his7-366 SH1L::ura4*$^+$ *SH2L::his7*$^+$ *SH2R::his7*$^+$ *SH3L::ura4*$^+$ |
| ST4932 (*972SD4[2L+]* #1) | *h*$^-$ *ura4-D18 SH1L::ura4*$^+$ *SH1R::his7*$^+$ *SH2R::his7*$^+$*SH3L::ura4*$^+$ |

ST5060 (*972SD4[2L+]* #2)

*h*$^{-}$ *ade6-M210 ura4-D18 SH1L*::*ura4*$^{+}$ *SH1R*::*his7*$^{+}$
*SH2R*::*his7*$^{+}$*SH3L*::*ura4*$^{+}$

ST5348 (*972SD4[2L+]* #3)

*h*$^{+}$ *ade6-M210 leu1-32 ura4-D18 his7-366 SH1L*::*ura4*$^{+}$ *SH1R*::*his7*$^{+}$
*SH2R*::*his7*$^{+}$*SH3L*::*ura4*$^{+}$

ST5349 (*972SD4[2R+]* #1)

*h*$^{-}$ *ade6-M216 leu1-32 ura4-D18 his7-366 SH1L*::*ura4*$^{+}$ *SH1R*::*his7*$^{+}$
*SH2L*::*his7*$^{+}$ *SH3L*::*ura4*$^{+}$

ST5350 (*972SD4[2R+]* #2)

*h*$^{+}$ *ade6-M210 leu1-32 ura4-D18 his7-366 SH1L*::*ura4*$^{+}$ *SH1R*::*his7*$^{+}$
*SH2L*::*his7*$^{+}$ *SH3L*::*ura4*$^{+}$

ST5351 (*972SD4[2R+]* #3)

*h*$^{-}$ *ade6-M210 SH1L*::*ura4*$^{+}$ *SH1R*::*his7*$^{+}$ *SH2L*::*his7*$^{+}$ *SH3L*::*ura4*$^{+}$

**Table S2. PCR primers used in this study.**

**<u>For amplification of the *SPBCPT2R1.03* ORF</u>**

| | |
|---|---|
| st498 | 5′- ATGAGTATTGAATTCGATGACAG-3′ |
| st499 | 5′- ACCATTCAAATTGTATTTCAAGTATG-3′ |

**<u>For amplification of telomere-distal *SH* regions</u>**

| | |
|---|---|
| ***subtel-5.2-7.9 kb*** | |
| jk618 | 5′-GCCTACCGCTTGCAGTTGTT-3′ |
| jk621 | 5′-ACAGTAAACTATGATCGCTTTTGAAGAC-3′ |
| ***subtel-7.9-9.3 kb*** | |
| jk620 | 5′-TTCTTAATCATTATCAAGTATTCATTGCAA-3′ |
| jk452 | 5′-CAACTTGCGAGAATGTAAACTACGTATTC-3′ |
| ***subtel-9.1-10.8 kb*** | |
| jk451 | 5′-TGCCTTTCTAGCCCATGACCTA-3′ |
| jk385 | 5′-CAGCATTAACCAACAGTGGTCTTC-3′ |
| ***subtel-10.7-13.2 kb*** | |
| jk384 | 5′-GCTCTCGACAAAGCCGTTCT-3′ |
| jk387 | 5′-CATCAGCAACGTCGCCAACT-3′ |
| ***subtel-13.0-15.1 kb*** | |
| jk386 | 5′-TTCCAAGTATGCCAGCTTATCATC-3′ |
| jk454 | 5′-TAAAGATTCATGGAACCATCATTTG-3′ |
| ***subtel-14.9-17.2 kb*** | |
| jk453 | 5′-AAAGAATTCGAATCACCCATACCA-3′ |
| jk683 | 5′-GGGTCAAAACCTGCGCATAA-3′ |
| ***subtel-17.1-19.0 kb*** | |
| jk682 | 5′-AATTTCAGTTGCCAAGGGACA-3′ |
| jk456 | 5′-GACCGCTACGCAACCATAAAG-3′ |
| ***subtel-18.9-21.3 kb*** | |
| jk455 | 5′-AACGAGTTGTGCAATGTTAGTAAGGT-3′ |
| jk561 | 5′-TGACCACCCTCATAAGCCTGT-3′ |
| ***subtel-21.3-23.3 kb*** | |
| jk560 | 5′-TGTTACGTGGCAAGGCAGACT-3′ |

jk559 5′-AACTCGCTTGATGAATGGGTG-3′

***subtel-23.3-26.2 kb***

jk558 5′-GACAACCCGACCCATGGAT-3′

jk555 5′-GACCCCAATGGCCGAGAT-3′

***subtel-26.2-27.7 kb***

jk554 5′-GCTGCATGTTTATATCCTTGGCT-3′

jk689 5′-GAGCCCAATGACCAAAATACAAA-3′

***subtel-27.7-29.0 kb***

jk688 5′-CAGCCCTGGGTGCAACTATAA-3′

jk389 5′-TGTTACTCATACTTGAAATACAATTTGAATG-3′

***subtel-34.9-36.9 kb***

tk74 5′-ACATTCAGAAACAACAGTTGG-3′

jk691 5′-GCTCCATCCATTGTCATTGGT-3′

**For detection of lengths of telomere-distal *SH* regions (Figure 4B)**

***subtel2R-4.2-11.1 kb*** **(a)**

tk121 5′-ATTCTTTTCACTTTCTTTTCCTCG-3′

tk66 5′-CCATGATGTCGTTACGGC-3′

***subtel2R-9.3-15.7 kb*** **(b)**

jk451 5′-TGCCTTTCTAGCCCATGACCTA-3′

tk79 5′-CATTCATTTTCTACACGCTCTATTATTTC-3′

***subtel2R-14.8-20.7 kb*** **(c)**

jk1611 5′-CATCCGGCGATATCTCTAGCAG-3′

tk44 5′-CAAAGTTGTATTCATGCTTGTTAC-3′

***subtel2R-20.6-22.1 kb*** **(d)**

tk114 5′-TCATTACCAAAGTTTAAAACGC-3′

tk113 5′-AGTCTGAATGTGCCAACTC-3′

***subtel2R-22.1-26.7 kb*** **(e)**

tk122 5′-GAGTTGGCACATTCAGACT-3′

tk123 5′-GGTATCGGTCACCGATATAGC-3′

***subtel2R-26.7-35.4 kb*** **(f)**

tk102 5′-GCTATATCGGTGACCGATACC-3′

tk130 5′-GGGAGTCCAAATTGGTTCAA-3′

***subtel2R-35.4-40.9 kb*** **(g)**

| | |
|---|---|
| tk76 | 5′-TTGAACCAATTTGGACTCCC-3′ |
| tk144 | 5′-GGTGTTGCGGGATCTTCA-3′ |

***subtel2R-40.7-44.1 kb*** **(h)**

| | |
|---|---|
| jk692 | 5′-CGCGCACTTTTCGGACATA-3′ |
| tk115 | 5′-GTATGATTTTTCAAACTGTCCC-3′ |

***subtel2R-44.1-47.5 kb*** **(i)**

| | |
|---|---|
| tk124 | 5′-GGGACAGTTTGAAAAATCATAC-3′ |
| tk125 | 5′-CAACGACTAAGTCTATACATCGG-3′ |

***rap1*** **(control)**

| | |
|---|---|
| jk900 | 5′-AATTGTAAAGCCTTACTCGGCCCAAAAGGATTATACACAA-AG-3′ |
| jk1219 | 5′-ATGAAGAATCTGGAGTTGACGTAG-3′ |

**For *Ch2R* sequencing**

***subtel2R-17.1-19.8 kb***

| | |
|---|---|
| jk682 | 5′-AATTTCAGTTGCCAAGGGACA-3′ |
| tk52 | 5′-GGAATTGATTATTCAACTCATTGGAC-3′ |

***subtel2R-29.0-30.9 kb***

| | |
|---|---|
| jk388 | 5′-GGCACAATTTCATTTCGTTTAGTTTAC-3′ |
| jk685 | 5′-TGCCGGTATTTAAGATGCACAT-3′ |

***subtel2R-34.6-36.9 kb***

| | |
|---|---|
| jk688 | 5′-CAGCCCTGGGTGCAACTATAA-3′ |
| jk691 | 5′-GCTCCATCCATTGTCATTGGT-3′ |

***subtel2R-50.8-52.3 kb***

| | |
|---|---|
| jk700 | 5′-TCCGCAAATTTTGTTAGCCAT-3′ |
| jk703 | 5′-ACGGGCGGTATGGATCAAT-3′ |

***subtel2R-54.8-57.0 kb***

| | |
|---|---|
| jk704 | 5′-TTCCAATCCCAGAGTCGTGTC-3′ |
| jk707 | 5′-AATAGACTTGGTCTGGATGAAGCAG-3′ |

***subtel2R-69.0-72.2 kb***

| | |
|---|---|
| jk710 | 5′-GCTTTTATGCAATGTCTTGACGA-3′ |
| tk131 | 5′-CCTTGATACATGGAACCTCG-3′ |

***subtel2R-83.6-86.4 kb***

| | |
|---|---|
| tk133 | 5′-AGTCCTTGTCCGGCAGAATCT-3′ |
| tk132 | 5′-AAGCTATCTGGGAAGCCG-3′ |

***subtel2R-98.0-102.7 kb***

| | |
|---|---|
| km38 | 5′-TTCAGGATTAAGGTAACGCGGT-3′ |
| st104 | 5′-TTTATGCATCCAACCGTTCGA-3′ |

***2R-106.2-110.0 kb***

| | |
|---|---|
| st214 | 5′-GCTTTACTCCTTTCAGGCGTATG-3′ |
| km41 | 5′-CGATCGTCCACTACTTCACGTTT-3′ |

***rap1***

| | |
|---|---|
| jk1065 | 5′-TATAAAGGATCCGTCATTTACATTCACCAAAAGCG-3′ |
| jk1219 | 5′-ATGAAGAATCTGGAGTTGACGTAG-3′ |

**For detection of RNA from upstream of the *tlh* ORFs (Figure 6C)**

| | |
|---|---|
| jk567 (a) | 5′-TTGCTGCTGCACAAAGCTTTTTG-3′ |
| tk79 (b) | 5′-CATTCATTTTCTACACGCTCTATTATTTC-3′ |
| jk1612 (c) | 5′-TGGTCGTCGCTTCAGAAATTG-3′ |
| tk41 | 5′-ATGTTTTGTGCATCTTCTTTCTCC-3′ |

***rap1* (control)**

| | |
|---|---|
| jk1300 | 5′-TATAAAGAATTCTCTAAATGAAGTTTCACGCAAGGT-3′ |
| jk1206 | 5′-TATAAACCCGGGGTATCCTCAACATTCTCTTGAATC-3′ |

**For detection of RNA from *dh*-homologous regions in the *tlh* genes (Figure 6D)**

***dh-homologous region***

| | |
|---|---|
| jk458 | 5′-TCGATAGAGATACGTTCATCTCATAGTGT-3′ |
| jk457 | 5′-CACTCATTTCTTTGCTGATCACTTACT-3′ |

***his1* (control)**

| | |
|---|---|
| jk1335 | 5′-CGAAGACGTGCTTCAGCGA-3′ |
| jk1336 | 5′-TGTCCACCTCGGAATCACTG-3′ |

**Table S3. Sequence identities between sequences of *2R* in *972SD4[2R+]* (#1 and #2) and those in PomBase**

| | Locus | Positions in PomBase (2R) | vs. PomBase *972SD4[2R+]* #1 Match / Total (bp) | vs. PomBase *972SD4[2R+]* #1 Identity (%) | vs. PomBase *972SD4[2R+]* #2 Match / Total (bp) | vs. PomBase *972SD4[2R+]* #2 Identity (%) |
|---|---|---|---|---|---|---|
| Subtelomere | *tlh2* | 4526885 - 4532644 | 7408 / 7452 | 99.41 | 7409 / 7454 | 99.40 |
| Subtelomere | Telomere-distal *SH* | 4521826 - 4524375 | 2549 / 2550 | 99.96 | 2549 / 2550 | 99.96 |
| Subtelomere | Telomere-distal *SH* | 4510792 - 4512641 | 1850 / 1850 | 100.00 | 1850 / 1850 | 100.00 |
| Subtelomere | Telomere-distal *SH* | 4504812 - 4506754 | 1943 / 1943 | 100.00 | 1943 / 1943 | 100.00 |
| Subtelomere | *SH*-adjacent | 4489185 - 4490784 | 1600 / 1600 | 100.00 | 1600 / 1600 | 100.00 |
| Subtelomere | *SH*-adjacent | 4484916 - 4486915 | 2000 / 2000 | 100.00 | 2000 / 2000 | 100.00 |
| Subtelomere | *SH*-adjacent | 4469756 - 4472805 | 3050 / 3050 | 100.00 | 3050 / 3050 | 100.00 |
| Subtelomere | *SH*-adjacent | 4454962 - 4458161 | 3200 / 3200 | 100.00 | 3200 / 3200 | 100.00 |
| Subtelomere | Boundary | 4439149 - 4443731 | 4583 / 4583 | 100.00 | 4583 / 4583 | 100.00 |
| | Subtelomere-adjacent | 4431856 - 4435505 | 3650 / 3650 | 100.00 | 3650 / 3650 | 100.00 |
| | *rap1* | 3098435 - 3101084 | 2650 / 2650 | 100.00 | 2650 / 2650 | 100.00 |
